# Metabolic dysregulation of the lysophospholipid/autotaxin axis in the chromosome 9p21 gene SNP rs10757274

**DOI:** 10.1101/789768

**Authors:** Sven W. Meckelmann, Jade I. Hawksworth, Daniel White, Robert Andrews, Patricia Rodrigues, Anne O’Connor, Jorge Alvarez-Jarreta, Victoria J. Tyrrell, Christine Hinz, You Zhou, Julie Williams, Maceler Aldrovandi, William J Watkins, Adam J Engler, David A. Slatter, Stuart M Allen, Jay Acharya, Jacquie Mitchell, Jackie Cooper, Junken Aoki, Kuniyuki Kano, Steve E. Humphries, Valerie B. O’Donnell

## Abstract

**Aims:** Common chromosome 9p21 SNPs increase coronary heart disease (CHD) risk, independent of “traditional lipid risk factors”. However, lipids comprise large numbers of structurally-related molecules not measured in traditional risk measurements, and many have inflammatory bioactivities. Here we applied lipidomic and genomic approaches to three model systems, to characterize lipid metabolic changes in common Chr9p21 SNPs which confer ∼30% elevated CHD risk associated with altered expression of *ANRIL*, a long ncRNA.

**Methods and Results:** Untargeted and targeted lipidomics was applied to plasma samples from Northwick Park Heart Study II (NPHSII) homozygotes for AA or GG in rs10757274. Elevated risk GG correlated with reduced lysophosphospholipids (lysoPLs), lysophosphatidic acids (lysoPA) and autotaxin (ATX). Five other risk SNPs did not show this phenotype. Correlation and network analysis showed that lysoPL-lysoPA interconversion was uncoupled from ATX in GG, indicating metabolic dysregulation. To identify candidate genes, transcriptomic data from shRNA downregulation of *ANRIL* in HEK293 cells was mined. Significantly-altered expression of several lysoPL/lysoPA metabolising enzymes was found (*MBOAT2, PLA2G4C, LPCAT2, ACSL6, PNPLA2, PLBD1, PLPP1, PLPP2* and *PLPPR2)*. Next, vascular smooth muscle cells differentiated from iPSCs of individuals homozygous for Chr9p21 risk SNPs were examined. Here, the presence of risk alleles was associated with altered expression of several lysoPL/lysoPA enzymes. Importantly, for several, deletion of the risk locus fully or partially reversed their expression to non-risk haplotype levels: *ACSL3, DGKA, PLA2G2A, LPCAT2, LPL, PLA2G3, PNPLA3, PLA2G12A LIPC, LCAT, PLA2G6, ACSL1, MBOAT2.*

**Conclusion:** A Chr9p21 risk SNP associates with complex alterations in immune-bioactive phospholipids and their enzymatic metabolism. Lipid metabolites and genomic pathways associated with CHD pathogenesis in Chr9p21 and *ANRIL*-associated disease are demonstrated.

**One sentence summary:** Inflammatory phospholipid metabolism defines a cardiovascular disease SNP

## Introduction

The association of altered plasma “lipids” with coronary heart disease (CHD) risk has been known for decades, however for some CHD-risk SNPs, there is no association with “traditional lipid measurements”, such as lipoproteins (HDL or LDL) or their constituents: cholesteryl esters (CE) and triglycerides (TG) ^1^. As a prominent example, the relatively common *CDKN2A/2B* (rs10757274, A>G) (minor allele frequency = 0.48) SNP on chromosome 9p21 confers ∼30% elevated risk of CHD, but acts independently of traditional lipid risk factors ^1^. Chr9p21 SNPs, including rs10757274, are believed to alter disease risk through modulation of the long non-coding (lnc)RNA, *ANRIL*, although both up and downregulation has been associated with risk ^*2, 3*^ The *ANRIL* product is detected in peripheral blood cells, aortic smooth muscle, endothelial cells and heart, and SNPs in Chr9p21 are associated not only with CHD but also numerous cancers ^2, 4-6^. Cellular studies show that *ANRIL* lncRNA down-regulates the tumour suppressors *CDKN2A/2B* by epigenetic regulation, modulating expression of pathways involved in differentiation, apoptosis, matrix remodelling, proliferation, apoptosis, senescence and inflammation ^5, 7^.

Lipids represent thousands of diverse molecules. However, CHD clinical risk algorithms such as Framingham or QRISK include circulating lipoproteins only ^8, 9^. Importantly, bioactive lipids that regulate vascular inflammation/proliferation in line with the function of *ANRIL* and thus maybe directly relevant to Chr9p21-medicated CHD are not included in these measures. Indeed, whether *ANRIL* mediates its effects via an impact on bioactive lipid signalling has not been examined, and was studied herein using lipidomics.

Here, plasma from a prospective cohort (Northwick Park Heart Study II, NPHSII) which recruited ∼3,000 men aged 50 - 61 years clinically free of CHD in 1990-1991, was analysed using targeted and untargeted lipidomics, followed by validation, metabolic correlation and network analysis ^10, 11^. Then, gene transcription for lipid metabolic enzymes was mined in data from a cellular *ANRIL* knockdown study, and from vascular smooth muscle cells differentiated from iPSCs obtained from individuals carrying Chr9p21 risk SNPs^12, 13^.

## Methods

### (more details methods are in Supplementary Data)

#### Patient samples

NPHSII is a prospective CHD study of ∼3000 men ^10, 11^. Middle-aged men (aged 50–64 yrs) were recruited from 9 general practices in the UK 27-yrs ago. Exclusions included a history of CHD or diabetes. Full information on the cohort is in Supplementary Methods. SNPs were chosen based on known association with altered risk of CHD and described in full in Supplementary Methods. Details of sample sizes, genes, SNPs and average levels of total triacylglycerides (TAG) and total cholesterol determined for these samples, are provided in Table 1. Samples were randomly chosen.

**Table 1:** Summary of genes, SNPs, sample numbers and recruitment lipid levels used in the untargeted analysis. ^$^ All subjects were homozygous for the common alleles for all other selected SNPs. * p < 0.05, ** p < 0.01, *** p < 0.005, 2-tailed, unpaired Student’s T-test, significantly different to APOE E3E3 controls.

| Gene | SNP | Genotype | n | total cholesterol<br>(mean ± SD) | total triglycerides<br>(mean ± SD) | CVD<br>incidence |
| --- | --- | --- | --- | --- | --- | --- |
| *APOE*<br>(controls) | E3 | E3E3<sup>§</sup> | 39 | 6.02 ± 0.96 | 1.91 ± 1.17 | 2 |
| *APOE* | E2 | E2E2 | 21 | 5.08 ± 0.99*** | 2.15 ± 0.99 | 0 |
| *APOE* | E4 | E4E4 | 37 | 5.69 ± 0.85 | 2.08 ± 1.39 | 4 |
| *APOA5* | rs662799 (A>G) | GG | 14 | 5.39 ± 0.82* | 2.76 ± 1.25* | 4 |
| *SORT1* | rs599839 (A>G) | GG | 38 | 5.45 ± 0.81** | 1.79 ± 0.87 | 5 |
| *LDLR* | rs6511720 (G>T) | TT | 21 | 5.21 ± 1.12** | 2.29 ± 2.16 | 4 |
| *CDKN2A/2B* | rs10757274(A>G) | GG | 33 | 5.64 ± 0.80* | 1.68 ± 0.80 | 6 |

#### Global lipidomics, informatics and statistics analysis

Lipids were extracted using two consecutive liquid-liquid extractions, first, hexane:isopropanol:acetic acid, then a modified Bligh and Dyer method as outlined in Supplementary Methods ^14^. Orbitrap datasets were processed using the R version of XCMS (Version 3.4), then using LipidFinder as described in Supplementary Methods ^15, 16^. This enabled assignment of a putative lipid class to 30 – 50 % of all ions detected. Our approach to statistical analysis of global datasets is described in Supplementary Methods.

#### Targeted analysis of lysoPLs

Lipids were extracted from plasma using the liquid:liquid extraction method outlined in Supplementary Methods. LC-MS/MS was performed on a Nexera liquid chromatography system (Shimadzu) coupled to API 4000 qTrap mass spectrometer (Sciex) as in Supplementary Methods.

#### Measurement of ATX

The ATX levels in the plasma were determined using a two-site immunoenzymatic assay with an ATX assay reagent equipped with an automated immunoassay analyzer, AIA-2000 (Tosoh, Tokyo, Japan) as previously described ^17^.

#### Measurements of LPA

Quantification of LPA was performed according to a previous method with minor modification ^18^. Briefly, plasma samples were mixed with a 10-fold volume of methanol containing internal standard (17:0-LPA), then sonicated. After centrifugation at 21,500×g, the supernatants were filtered and subjected to LC-MS/MS. This consisted of a Ultimate3000 HPLC and a TSQ Quantiva triple quadropole mass spectrometer (Thermo Fisher Scientific, San Jose, CA).

#### Analysis of Affymetrix data from ANRIL down-regulation in cell lines and RNAseq data from iPSC-derived VSMCs

Raw Affymetrix CEL files relating to total transcript expression in HEK 293 cells stimulated with Tetracycline (shRNA ANRIL silenced for 0h, 48h and 96h) were downloaded from the GEO database (accession: GSE111843) and analysed using packages in CRAN and Bioconductor: limma, oligo, ggplot2 ^12, 19-22^, as described in Supplementary Methods. Data from iPSC-derived VSMCs available at GEO (GSE120099) was analysed as described in Supplementary Methods.

#### Statistics

Statistics for untargeted lipidomics was performed as described in Supplementary Methods. Targeted data are shown as Tukey box plots, * p < 0.05, ** p < 0.01, *** p < 0.005, Mann Whitney U and Student’s t-test were used as described in legends. Correlation analysis was undertaken using Answerminer (https://www.answerminer.com/calculators/correlation-test), using Pearson’s correlation coefficient. To compare the slopes (or “Pearson correlation coefficients”) of regression lines between AA and GG carriers, we used the method described ^23^. See Supplementary Methods for more details on statistical methods used.

## Results

### Global lipidomics demonstrates that lysoPLs are reduced in GG plasma versus AA

To capture all lipids (knowns/unknowns), high resolution Orbitrap MS data from long chromatographic separations was analysed using XCMS, then processed for cleanup/assignment to LIPID MAPS categories, using LipidFinder (Figure 1 A,B) ^16^. Plasma quality was checked through careful comparison with fresh plasma, detailed in Supplementary Data. Most lipid categories were unchanged, however oxidized phospholipids and lysoPCs had elevated somewhat in storage (Supplementary Figures 1-3). This is not unexpected and we include a full discussion of this phenomenon in Supplementary Data. To assess the impact of the rs10757274, A>G SNP, we compared AA (n = 39) with the risk genotype GG (n = 33). The full dataset is provided as Supplementary Data (supplementary data.xlsx, tab 1). Data was analysed first using a Mann Whitney U test, then chromatograms for all features with p<0.075 were manually checked for quality. LipidFinder detected 1878 lipids, with 872 assigned to a category (Figure 1 B). Next, quantile normalization was applied followed by Mann Whitney U test, and then a p-value adjustment using sequential goodness of fit metatest (SGoF) to each subclass^24^. The SGoF has been shown as especially well suited to small sample sizes when the number of tests is large. This data is shown in volcano plots in Figure 1 C-J, and the p-values are in column M (Supplementary Data.xls, tabs 1,2). Those most affected by genotype were GPLs and unknowns (Figure 1 C-J, Table 2). Following p-value adjustment the number of significantly different lipids was 17, with 7 putatively identified as lysoPC ions and adducts (supplementary data.xlsx, tab 1,2). An additional group of 8 had p-values close to significance at 0.05-0.08. All were reduced in GG plasma. As this method is used as for hypothesis generation only, we next validated our results using gold-standard quantitative targeted methods.

**Figure 1.**
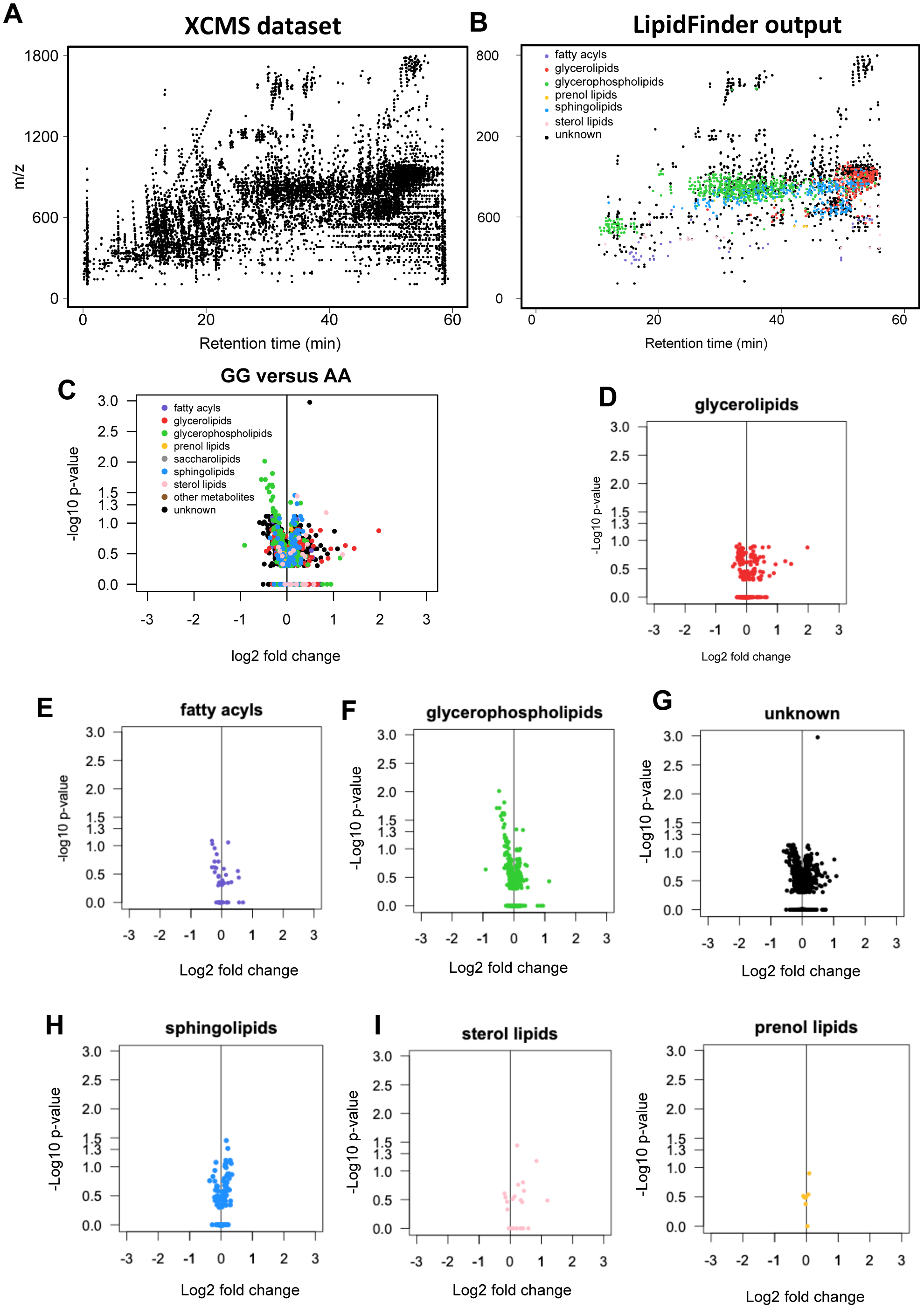
Global lipidomics reveals class specific changes in GPLs in rs10757274 GG vs AA. *Panel A: Scatterplot of features in a plasma sample (*≈*14,000) after processing high-resolution MS data using XCMS.* Analysis was undertaken using parameters provided in Supplementary Methods. *Panel B. Scatterplot obtained after LipidFinder and manual-data clean-up, as described in Methods*. Each dot represents a lipid described by *m/z* value and retention time. Putative identification and assignment of category was performed using WebSearch of the curated LIPID MAPS database. *Panels C-J. Volcano plots show differences in lipid classes with genotype.* Volcano plots were generated as described in Methods, plotting log2(fold-change) versus – log10(p-value) for all (n = 39 AA, 33 GG), following p-value adjustment using sequential goodness of fit metatest (SGoF).

**Table 2:** Number of detected and identified lipid features in the global lipidomics assay. The number of lipids in each class are shown, with the number of significantly different lipids (non-parametric one-tailed Mann–Whitney U test, assuming unequal variance with a threshold of p ≤0.05, after SGoF correction), between the SNP group and controls shown in parentheses. FA: Fatty acyl, GPL: glycerophospholipid, GL: glycerolipid, SL: sphingolipid.

| Lipid Class | FA | GPL | GL | SL | Sterol | Prenol | Unknowns | TOTAL |
| --- | --- | --- | --- | --- | --- | --- | --- | --- |
| Detected<br>($P < 0.05$ after SGoF) | 51 (0) | 220 (0) | 401 (13) | 166 (2) | 27 (1) | 7 (0) | 1006 (1) | 1878 (17) |

### Quantitative targeted lipidomics confirms decreased lysoPLs in the GG samples

The same plasmas were analysed using a targeted fully-quantitative assay for 15 lysoPLs. Of these several lysoPCs significantly decreased, with both lysoPC and lysoPEs all trending towards lower levels in GG (Supplementary Figure 4 A). This was replicated using new samples from NPHSII (n = 47: AA, 49: GG) (Supplementary Figure 4 B). When both datasets were combined (n = 82 – 86/group), all 8 lysoPCs were significantly lower in the GG genotype (Figure 2 A). Thus, lysoPLs are overall suppressed in the GG genotype, with a more robust effect on lysoPCs than lysoPEs.

**Figure 2.**
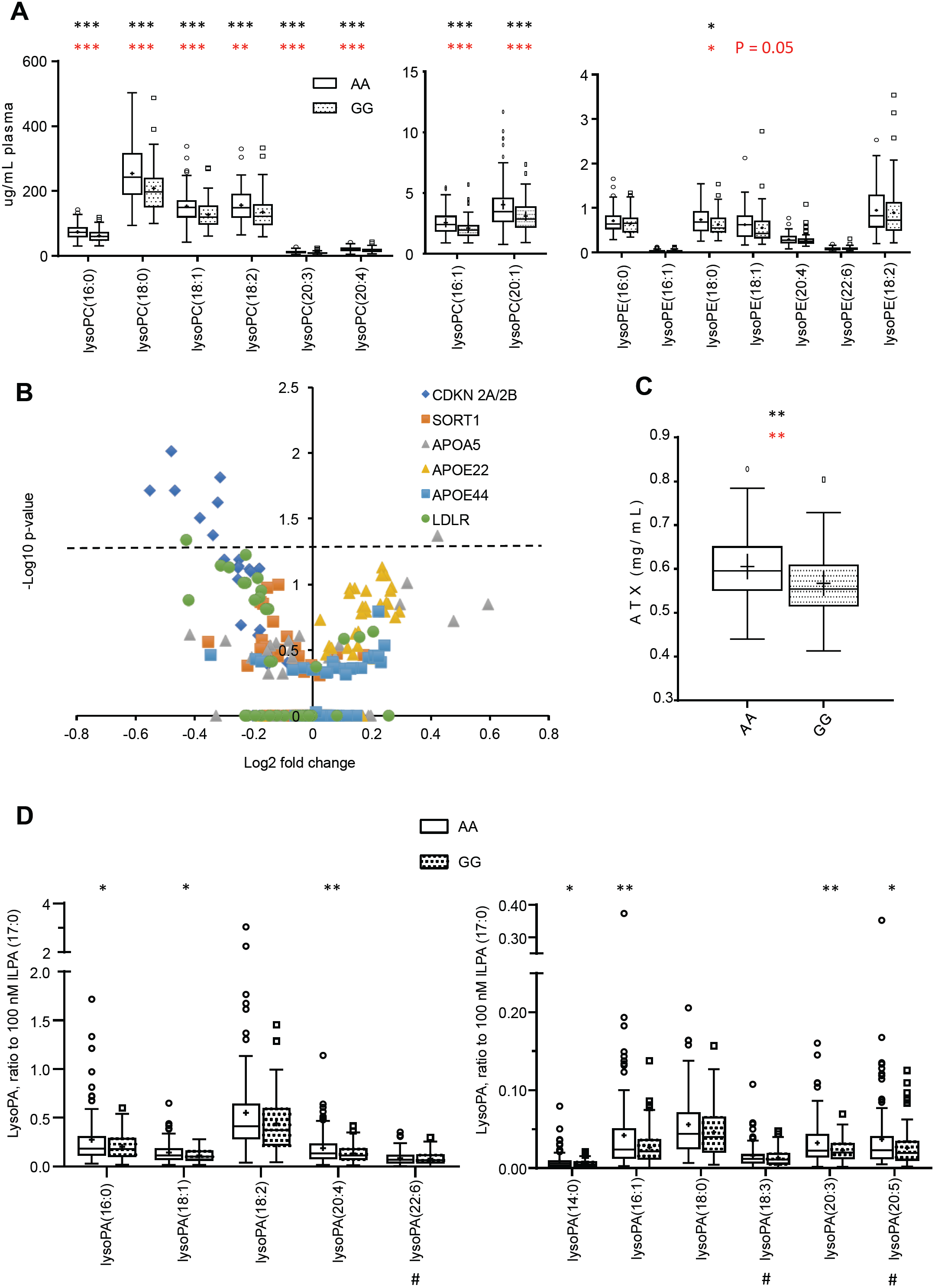
LysoPLs are significantly reduced in rs10757274 GG, but not in subjects with unrelated SNPs. *Panel A. Several LPCs are lower in GG samples than AA controls, and LPEs trend towards lower levels.* LysoPLs were determined using LC/MS/MS as described in Methods (n = 88 AA, 81 GG). Tukey box plot, * p < 0.05, ** p < 0.01, *** p < 0.005, 2-tailed, unpaired Student’s T-test (black) and Mann Whitney U (red). *Panel B. Plasma lysoPL are not altered by other risk SNPs.* Plasma from the NPHSII cohort containing several risk (up or down) SNPs were analysed using LipidFinder, and *m/z* values corresponding to lysoPL extracted and compared. These are plotted on a volcano plot, to show fold change *vs* significance, following p-value adjustment using sequential goodness of fit metatest (SGoF). Numbers and genotypes are shown in Table 1. *Panel C. ATX is significantly decreased in GG samples compared to AA controls.* Plasma ATX activity was measured as described in Methods (n = 47 AA, 49 GG). *Panel D. LysoPAs are significantly decreased in GG plasma compared to AA controls.* Plasma lysoPAs were measured as described in Methods, using LC/MS/MS (n = 95 AA, 100 GG). Tukey box plot, * p < 0.05, ** p < 0.01, *** p < 0.005, 2-tailed, unpaired Student’s T-test (black).

### Significantly altered lysoPLs are not detected in five additional CHD risk-altering SNPs

LipidFinder data was analysed for additional SNPs from the NPHSII cohort, comparing subjects homozygous for the common alleles with subjects homozygous for rare protective alleles for *SORT1, LDLR* or *APOE* E2/E2, or rare risk alleles *APOA5* or *APOE4/E4* (Table 1). For most lysoPLs, levels were not significantly altered, with the exception of one for *APOA5* (upregulated, lysoPE(18:1), and one for *LDLR* (downregulated, lysoPC(18:2)) (Figure 2 B). This indicates that lysoPLs are consistently reduced only in the GG risk SNP rs10757274.

### The plasma lysophosphatidic acids(lysoPA)/autotaxin (ATX) axis is dysregulated in the GG group

Next, lysoPL-related metabolites/enzymes were measured. Metabolism of lysoPL to lysoPA in healthy plasma can be mediated by ATX^25^. Here, we used a targeted LC/MS/MS assay for lysoPAs, and an immunoenzymatic assay for ATX. ATX was significantly decreased (p = 0.026). Based on power calculations (Supplementary Data), an additional set of plasmas was included to increase sample numbers to 95-100 per group for lysoPAs. LC/MS/MS demonstrated overall small reductions, but with several being significantly lower (Figure 2 C,D). Taken with the lysoPL data, this indicates a global suppression of lysoPL/lysoPA/ATX metabolic pathway in the GG group.

Next, correlation analysis was undertaken to determine the contribution of ATX in metabolizing lysoPL to lysoPA. In AA plasmas, ATX showed weakly negative or positive correlations with total lysoPL or lysoPA, respectively (Figure 3 A,B). This agrees with reports that ATX contributes to lysoPL conversion to lysoPA in healthy subjects ^25^. In contrast, in GG plasmas, these trends were reversed (Figure 3 C,D). Next, we correlated substrates with products (Figure 3 E-H). In the AA group, significant positive correlations were seen for total lysoPA with lysoPL (p = 0.034). Comparing lipids with the same fatty acyl, significant correlation was seen between lysoPA(18:2) and lysoPL(18:2) (p = 0.023) (Figure 3 E,F). This indicates that as the pool of lysoPL increases, the level of lysoPA increases in parallel, consistent with conversion by ATX. This relationship was fully reversed in the GG group, where total lysoPL, lysoPL(18:2) or lysoPL(20:4) were negatively correlated with their corresponding lysoPAs (p = 0.019, 0.054, 0.019 respectively) (Figure 3 G-I). We next analysed correlation slopes for AA *versus* GG, comparing either lysoPL:lysoPA (Figure 3 E *versus* G), or lysoPL(18:2):lysoPA(18:2) (Figure 3 F *versus* H). Both these comparisons revealed significant differences (p = 0.0264 and 0.0029 respectively) ^23^. These data confirm altered metabolism of lysoPL and lysoPA lipids between genotypes. Specifically, conversion of lysoPL to lysoPA is suppressed in the GG homozygotes.

**Figure 3.**
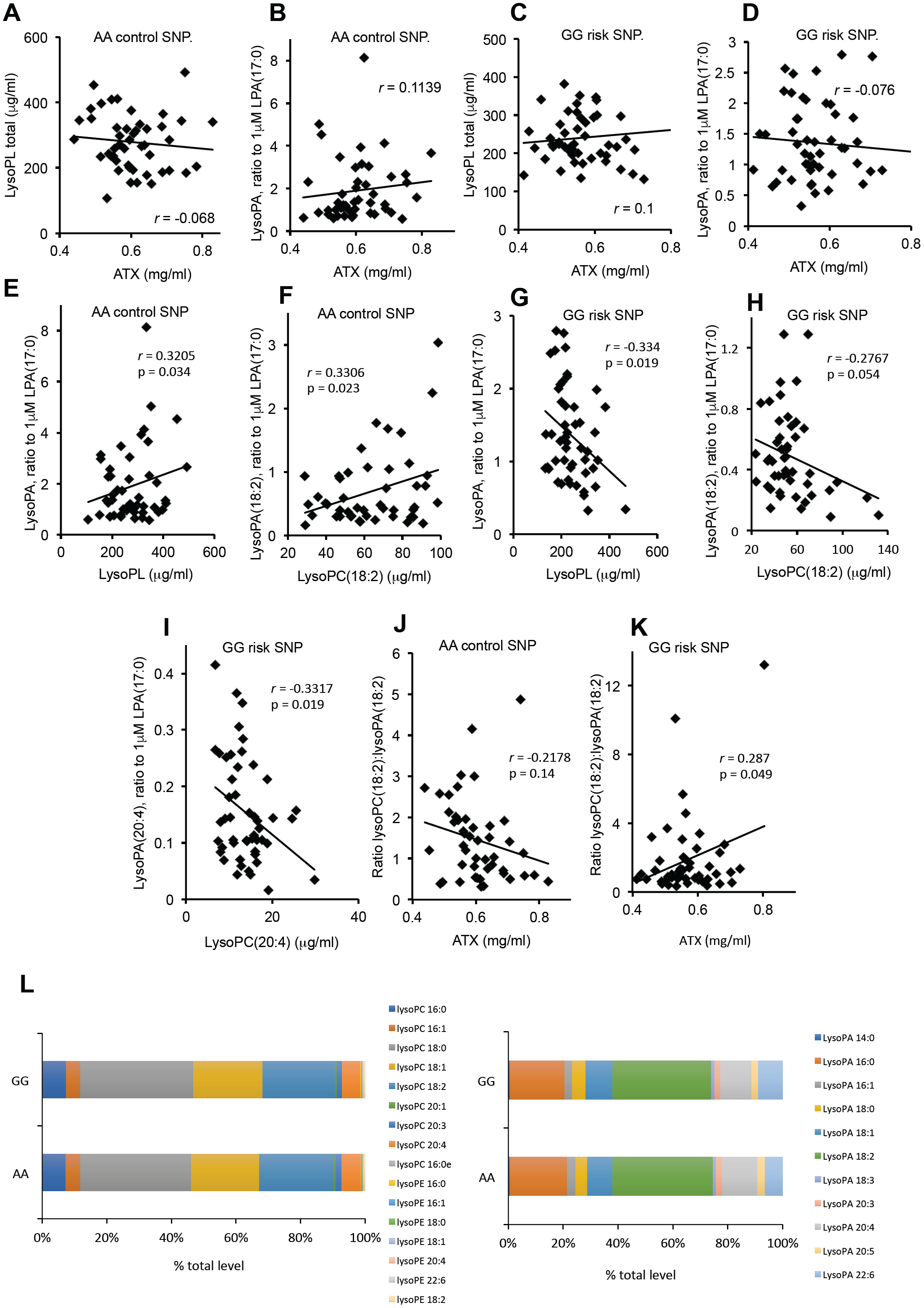
The lysoPL/lysoPA/ATX axis is dysregulated in the GG plasmas, while the profile of molecular species is unchanged for lysoPL/lysoPA. *Panels A-D. ATX shows altered correlations with plasma lysoPL or lysoPA in GG versus AA plasma*. Levels of lysoPL or lysoPA quantified by LC/MS/MS in the validation cohort were correlated using Answerminer, to determine Pearson’s correlation co-efficient. *A,B*: AA control plasma, *C,D*: GG risk plasma (n = 47 AA, 49 GG). *Panels E-I. LysoPL and lysoPA are positively correlated for AA plasma, but negatively correlated for GG*. The sum of all lysoPAs or lysoPLs in each set were correlated using Answerminer, as above (E,G). Alternatively, lipids containing 18:2, or 20:4 were separately correlated (F,H,I). *E,F*: AA control plasma, G,H,I: GG risk plasma (n = 47 AA, 49 GG). *Panels J,K. The lysoPA(18:2)/lysoPL(18:2) ratio positively correlates with ATX in AA plasma, but negatively for GG, indicating a block in substrate:product conversion in GG*. Correlations were performed using Answerminer (n = 47 AA, 49 GG). *J*: AA plasma, *K*: GG plasma. P<0.05 indicates significant using Pearson’s correlation test. *Panel L. The profile of individual lysoPL or lysoPA molecular species is unchanged between GG and AA plasmas*. Levels of individual lysoPL/lysoPA were compared across both groups, and shown as %.

The direct contribution of ATX to metabolizing lysoPL to lysoPA was next examined by correlating normalized ratios of lysoPC(18:2):lysoPA(18:2) with ATX. In this comparison, we expect that as ATX increases, the ratio of substrate:product will reduce due to their interconversion. Indeed, the For AA plasma, a weak negative correlation was seen (Figure 3 J). In contrast, a significant positive correlation was observed for GG plasma (Figure 4 K). Thus, as ATX increases, a higher ratio of substrate:product was seen in GG, decoupling ATX from metabolizing lysoPL to lysoPA. Comparing the slopes for AA *versus* GG revealed significant differences based on genotype (p = 0.0157). This further underscores the dysregulation of the lysoPL metabolic pathway in the GG group, suggesting that non-ATX pathways mediate lysoPL to lysoPA conversion. Last, the relative ratios of all lysoPL and lysoPA molecular species were unchanged in the GG versus AA groups (Figure 3 L). Thus, while metabolism of lysoPL/lysoPA by ATX is altered, there was no influence of genotype on molecular composition overall.

**Figure 4.**
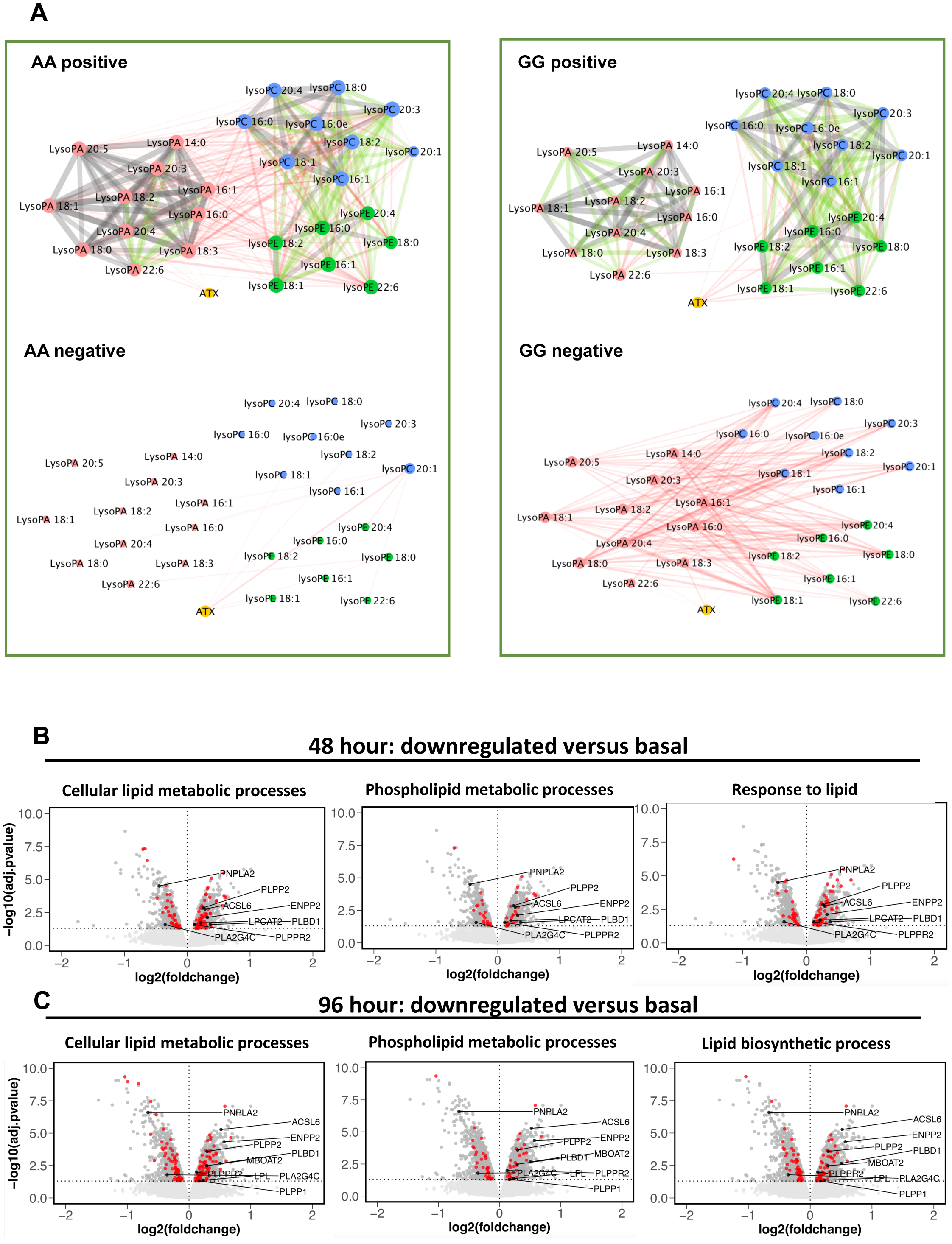
Cytoscape analysis of lipids reveals divergent metabolism in GG *versus* AA, while *ANRIL* knockdown is associated with significant changes to lysoPL/lysoPA-metabolising genes. *Panel A. Cytoscape reveals strong links within related families, but a positive-negative switch for lysoPL-lysoPA correlations between AA-GG plasmas.* Pearson correlation networks were generated for the AA and GG validation samples (n = 47 AA, 49 GG), using lipid concentrations. Nodes are coloured by lipid sub-category and represent individual molecular species, and edges represent the correlation. Edges detail the Pearson correlation coefficients between nodes (lipids), where the width of the edge denotes value. Additionally, edges are coloured by value: red (r = 0.10-0.39); green (r = 0.40-0.69); grey (r 0.70-1.00). *Panels C,D. Significant changes in lipid regulatory gene expression are observed with ANRIL knockdown in cell culture.* Affymetrix array data generated in ^5^ was analysed using GO as described in Methods. Volcano plot showing differential gene expression of all genes on the Affymetrix HuGene1.0 v1, chip. LysoPL/lysoPA regulating genes that alter in line with decreased levels of the lipids in GG plasma are labelled. The horizontal dashed line shows where adj.pvalue < 0.05 (Benjamini-Hochberg correction) where points (genes) above this line are significantly differentially expressed. LysoPL-regulating genes that alter in line with decreased levels of the lipids are labelled in black. Genes in red are annotated to the GO-term detailed in the plot title. Data are plotted in R using ggplot2. *Panel B*: 48-hr shRNA knockdown, *Panel C*: 96-hr shRNA knockdown.

Next, a Pearson correlation analysis looking at relationships between individual lipids and ATX was next undertaken using Cytoscape. For thresholds, the classification system of Schober was used ^26^. Here, we see that there are moderate (r = 0.40-0.69, green) or strong (r = 0.70-1.00 grey) correlations between lipids of the same class, while there are weak (r = 0.10-0.39, red) correlations between different lipid classes (Figure 4 A). Importantly, the key difference in the dataset is that the weak correlations between classes are positive for the AA group, while they are negative for the GG group (Figure 4 A). Overall, this indicates that these lipids behave similarly within AA subjects. In contrast, in GG plasma, while lipid classes still positively correlate within their groups (e.g. lysoPCs correlate strongly with each other), the links between lysoPL and lysoPA are lost. Instead correlations were negative between lysoPE and lysoPA (Figure 4 A). As in Figure 4, ATX weakly positively correlates with lysoPA in the AA group, but instead with lysoPL in the GG group. This analysis reinforces our findings of altered metabolism for lysoPL/lysoPA, but here at the level of individual lipid species.

### ANRIL knockdown significantly alters lipid and lysoPL metabolism gene expression

Chr9p21 risk SNPs are believed to act via altering expression of *ANRIL*, which regulates cell proliferation/senescence *in vitro* ^2, 4, 5^. To examine for a functional link with lysoPL/lysoPA metabolism, we analysed the effect of shRNA downregulation of the proximal *ANRIL* transcripts EU741058 and DQ485454 in HEK 293 cells at 48 hrs and 96 hrs ^12^. A GO analysis found significant alterations of several lipid pathways by *ANRIL*, including *Regulation of Lipid Metabolic Processes* (GO: 0019216), *Phospholipid Metabolic Processes* (GO:0006644), *Cellular Lipid Metabolic Process* (GO:0044255) and *Lipid Biosynthetic Processes* (GO:0008610), for example, *Regulation of Lipid Metabolic Processes* was 1.9 or 1.88 fold-enriched (FDR < 0.05, Benjamini-Hochberg) respectively at 48 and 96-hrs respectively (Table 3, Supplementary Data.xlsx, tabs 3,4). Thus, large numbers of lipid-associated genes were significantly differentially regulated (Supplementary Data.xls, tabs 5,6).

**Table 3.** Several lipid related Gene Ontology Pathways are significantly regulated by ANRIL silencing in HEK 293 cells. Results are from the PANTHER Over-representation Test, Term enrichment service (pantherdb.org), using default analysis parameters (Fisher’s Exact test with False Discovery Rate: FDR P < 0.05, Benjamini-Hochberg). Fold-enrichment represents the number of observed differentially expressed genes with the GO annotation of interest, relative to genome background. The full list of genes significantly altered in these GO processes, is provided in Supplementary Data.xlsx, along with a list of all significantly-altered GO processes at both timepoints.

|  | GO biological process | Significantly different genes | Fold enrichment | P-value | FDR |
| --- | --- | --- | --- | --- | --- |
| **48 hrs shRNA knockdown versus control** | glycosphingolipid metabolic process (GO:0006687) | 14 | 3.05 | 5.68E-04 | 3.12E-02 |
|  | regulation of lipid metabolic process (GO:0019216) | 46 | 1.9 | 1.12E-04 | 9.22E-03 |
|  | phospholipid metabolic process (GO:0006644) | 51 | 1.85 | 9.16E-05 | 7.99E-03 |
|  | cellular lipid metabolic process (GO:0044255) | 109 | 1.68 | 6.07E-07 | 1.37E-04 |
|  | lipid metabolic process (GO:0006629) | 130 | 1.61 | 5.02E-07 | 1.19E-04 |
|  | response to lipid (GO:0033993) | 84 | 1.53 | 2.59E-04 | 1.74E-02 |
| **96 hrs shRNA knockdown versus control** | regulation of lipid metabolic process (GO:0019216) | 74 | 1.88 | 2.80E-06 | 3.38E-04 |
|  | phospholipid metabolic process (GO:0006644) | 73 | 1.63 | 2.05E-04 | 1.30E-02 |
|  | lipid biosynthetic process (GO:0008610) | 102 | 1.61 | 1.54E-05 | 1.51E-03 |
|  | cellular lipid metabolic process (GO:0044255) | 164 | 1.56 | 2.74E-07 | 4.70E-05 |
|  | lipid metabolic process (GO:0006629) | 203 | 1.54 | 1.61E-08 | 3.79E-06 |

We next examined the effect of *ANRIL* knockdown on 48 candidate lysoPL metabolism genes (Supplementary Data.xlsx, tab 7). Of these, 9 were significantly changed at both timepoints, and another 6 at a single timepoint (Table 4). Several were consistent with lowered lysoPL/lysoPA including reduced *PNPLA2, PLA2G4C*, increased *LPCAT2, MBOAT2, ACSL6, PLBD1, PLPP1, PLPP2* and *PLPPR2* (Table 4, Scheme 1). Additional relevant genes were regulated, but in the opposing direction, including decreased *LPCAT1* and *LPCAT3* and increased *LPL, PLA2G7*, and *DGKA* (Table 4). *ENPP2* (the gene encoding ATX) was significantly increased by *ANRIL* suppression (Table 4, Scheme 1). This data is displayed in volcano plots of the full Affymetrix dataset (Figure 4 B,C, Supplementary Figure 5). Genes in red represent significantly-different lysoPL metabolizing genes from the lipid GO pathways.

**Table 4.**
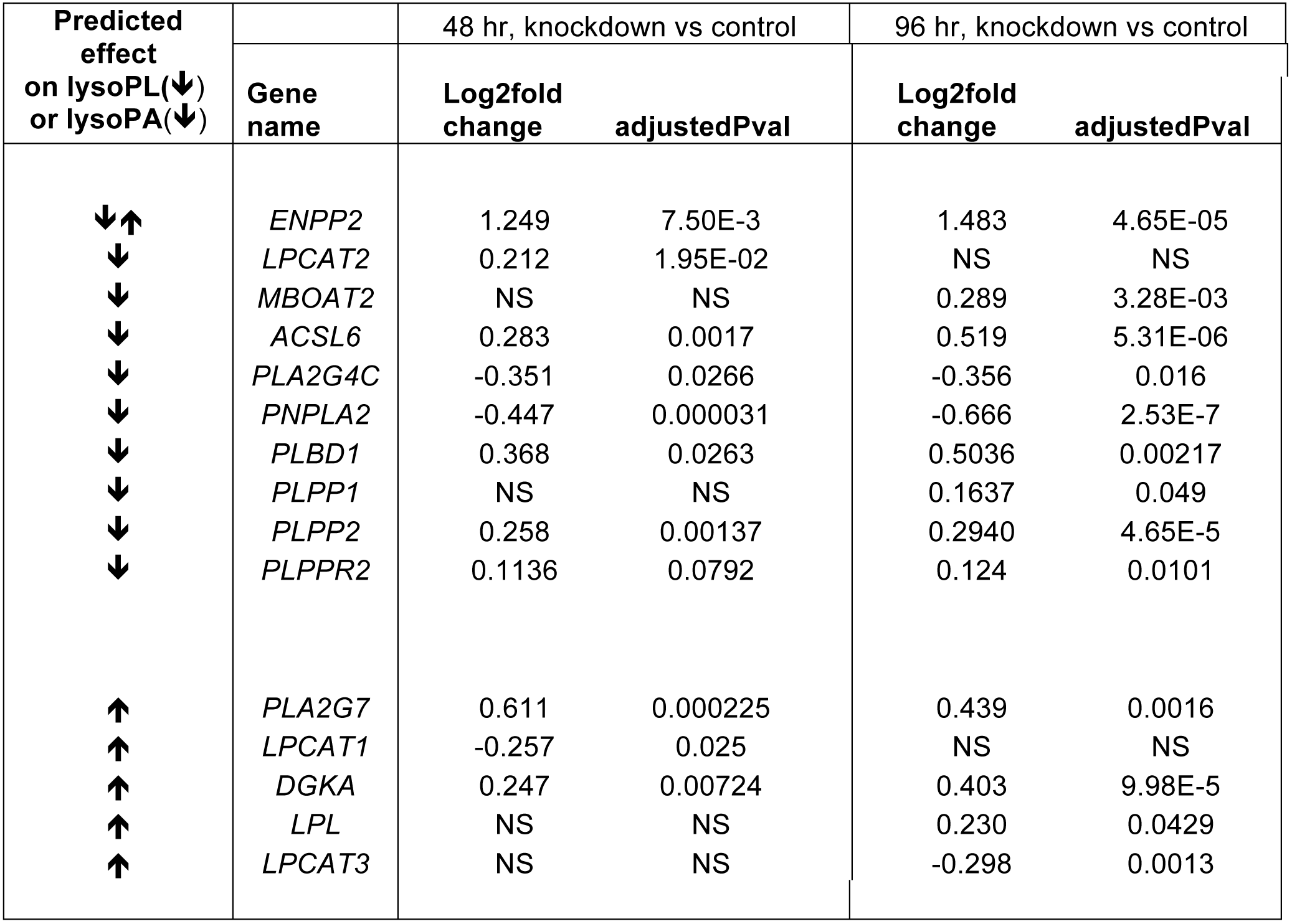
Several LysoPL relevant genes are significantly-altered in ANRIL knockdown. Data were analysed using the oligo and limma packages in Bioconductor, see methods. P values were corrected for multiple testing using Benjamini-Hochberg (adjusted p-value cut-off: 0.05)

### VSMCs generated from iPSCs from Chr9p21 risk haplotypes show altered expression of lysoPL metabolism genes

VSMCs generated by differentiation of iPSCs from humans homozygous for risk haplotypes in Chr9p21 show globally altered transcriptional networks, dysregulated adhesion, contraction and proliferation, with deletion of the risk haplotype rescuing the phenotype ^13^. Here, we interrogated the RNAseq dataset of mature iPSC derived VSMCs for expression of the same 48 lysoPL metabolism genes. Multivariate analysis using PCA shows clear separation of cell lines containing the risk haplotypes (RRWT) from controls (NNWT) in PC1 (Figure 5 A). When the risk locus was deleted, the resulting RRKO cell lines instead clustered with NNWT and NNKO (Figure 5 A). This indicates that overall expression of lysoPL metabolizing genes is different in risk haplotype cells, but reverts closer to non-risk (NN) on removal of the 9p21 locus. Examination of individual genes revealed 13 that were significantly different between NNWT and RRWT, where removal of the risk locus in RR led to partial or complete rescue: *ACSL3, DGKA, PLA2G2A, LPCAT2, LPL, PLA2G3, PNPLA3, PLA2G12A LIPC, LCAT, PLA2G6, ACSL1, MBOAT2* (Figure 5 B, Supplementary Figure 6).

**Figure 5.**
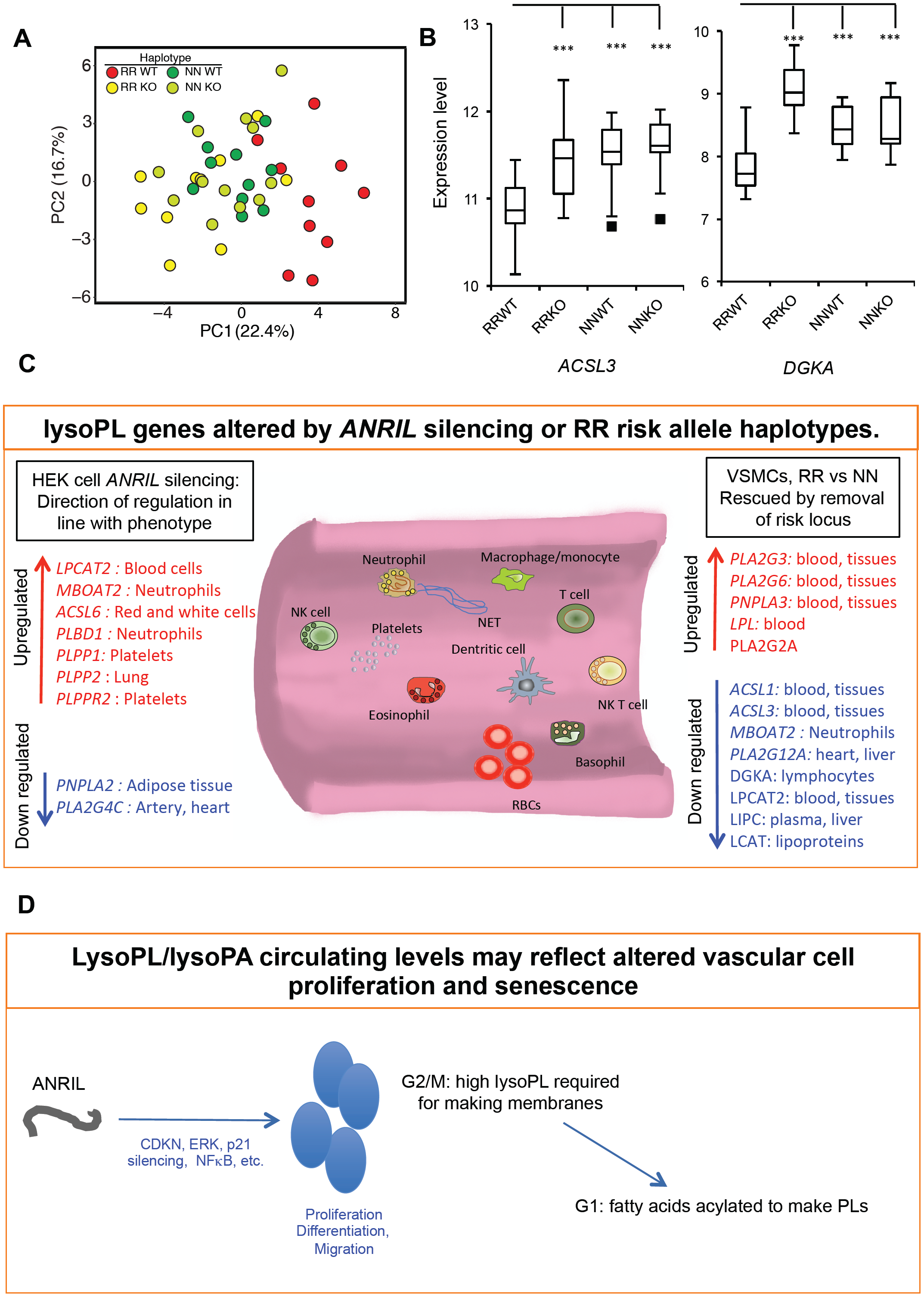
VSMCs from risk haplotypes show differential gene expression of lysoPL metabolizing genes, that are rescued by deletion of the Chr9p21 locus. *Panel A. PCA shows that the presence of risk haplotypes is associated with differential gene expression of lysoPL genes.* iPSCs from peripheral monocytes were obtained and differentiated as described in Supplementary Methods. RNAseq data was clustered using lysoPL metabolizing genes by PCA in R. Non-risk haplotype (NNWT), risk haplotype (RRWT) and their genome edited counterparts (NNKO and RRKO) are shown. *Panel B. Example datasets for ACSL3 and DGKA, showing that removing the risk locus reverts gene expression back to levels in non-risk individuals*. * p < 0.05, ** p < 0.01, *** p < 0.005, Students t-test. *Panels C,D. Schematics showing impact of ANRIL silencing or risk haplotypes on relevant lysoPL metabolizing genes.*

## Discussion

Lipidomics MS has been commonly applied to prospective CHD cohorts that contain no genetic information, while conversely GWAS studies have examined associations with traditional “lipid” measures only (e.g. total cholesterol or triglycerides) ^27-41^. Cohorts are only now starting to examine the association of individual lipid molecular species with specific risk SNPs, and little information on this is yet available. Here, we show that a common Chr9p21 (rs10757274, A>G) CHD-risk SNP is associated with metabolic alterations to the lysoPL/lysoPA/ATX axis (Figures 1-4). This revealed a genotype-specific defect absent in five other GWAS-proven CHD-risk SNPs. Since the action of rs10757274 GG is independent from “traditional lipid” measurements, it represents a different component of the disease, characterized by changes to bioactive signalling phospholipids (PL), rather than circulating storage/energy lipid pools ^1^. The three nearest genes to chromosome 9p21 are two cyclin-dependent kinase inhibitors *CDKN2A* and *CDKN2b*, and the non-coding RNA *ANRIL*. Compared to AA individuals, GG have almost 50% lower *ANRIL* transcription in peripheral blood cells ^2, 4, 5^. However, there are multiple *ANRIL* isoforms, including short, long and circular, and reports differ on which are specifically increased or decreased in CVD. In this regard, iPSC-derived VSMC lines from humans with risk haplotypes express higher levels of short ANRIL transcripts ^13^. The *ANRIL* gene product upregulates metabolic genes in cultured cells, as well as stimulating VSMC proliferation ^2, 5, 12^. Here, we showed that either shRNA knockdown of two proximal *ANRIL* transcripts or the presence of risk haplotypes in VSMCs from humans with risk haplotypes lead to multiple changes in PL-related lipid-metabolizing genes (Figure 4, Scheme 1). In the case of VSMCs, removal of the risk locus indicated that this was controlled by the Chr9p21 locus itself (Figure 5 A,B). Collectively, this suggests that Chr9p21 risk alleles may alter lysoPL/lysoPA in humans via *ANRIL* regulation, providing novel insights into the biology of this important cause of CHD. With the exception of *MBOAT2*, which was consistently elevated when *ANRIL* was lower, genes that were significantly altered were different between the *in vitro* datasets. This is likely a reflection of the complex and incompletely understood role of *ANRIL* in CVD risk, in line with the conflicting reports on its expression in risk haplotypes and its incompletely characterized multiple isoforms.

Our initial plasma screen comprised a relatively small sample size, while measuring large numbers of “features”. However, multiple comparison testing/correction (as in genomics), is problematic for this type of data for many reasons including: (i) multiple variables can represent the same metabolite, because it is impossible to remove all duplicates ions, (ii) lipidomic/metabolomics datasets are considerably more heterogeneous than genomics, at least in part because co-efficients of variation for MS data are relatively high. Here, we used LipidFinder as a screening tool only, to identify candidate lipids that we then validated using all the approaches outlined above. Next, we applied quantile normalization followed by Mann Whitney U test, and then a p-value adjustment using sequential goodness of fit metatest (SGoF) to each subclass. The SGoF has been shown to be especially well suited to small sample sizes when the number of tests is large^24^. This provides the level of confirmation required for identifying underlying pathways and mechanisms (rather than biomarkers). A similar approach led to the discovery (repeated now by many others) of the high importance of trimethylamine *N*-oxide (TMAO) in CHD pathogenesis ^42^.

Cohort lipidomics can be fraught with pitfalls, and it is critical these are taken into account to enable findings to be interpreted rigorously. Untargeted methods enable large numbers of samples to be analysed, but do not provide quantitation or full identification of lipids. This major issue inhibits cross-cohort data comparisons, and lacks validation. On the other hand, gold standard targeted methods are not generally amenable to large-scale screening. Here, we used untargeted lipidomics as a first screen only, avoiding naming lipids. This hypothesis-generating tool preceded a full replication and validation in the same and additional samples proving the observations were real, not false positives. We also combined two separate sample sets in our targeted approach to increase statistical power, as described^43^. LysoPCs/PAs can be formed/metabolized by enzymes in plasma, and the rate of this when samples are at room temperature is not trivial^44, 45^. This means that sample processing and storage are critical, and that it is essential that time-to-freezing is consistent. If not, it will impossible to reliably compare across samples both within and between cohorts. Lack of consistent sample processing is a major issue underlying well-known problems of reproducibility across cohort studies. In NPHSII, and the Bruneck cohort, which reported similar findings of lower lysoPC in subjects who subsequently developed CVD, sample processing was on site and fast, with immediate low temperature freezing following plasma isolation (Manuel Mayr, personal communication). Thus, artefactual generation of these lipids during plasma isolation will have been minimized, and the reduced levels found will either have occurred in vivo, or gradually during long term low temperature storage. Regardless, it is seen that AA and GG plasmas metabolize lysoPL/lysoPA differently, and understanding the underlying biological differences is important. We provide an expanded discussion on lipidomics methodologies and storage considerations in our Supplementary Data file.

LysoPL/lysoPAs were significantly lower in GG plasmas (Figure 2 A,D), and correlation of lipids and subjects revealed that their metabolism was altered in the risk SNP group (Figures 3, 4 A). This finding could indicate reduced involvement of ATX in metabolising lysoPL to lysoPA (consistent with the reduced levels of ATX detected), but it is likely that there are additional factors responsible that are not understood. One interpretation is that as ATX levels increase in this group, another metabolic pathway that either generates lysoPC or removes lysoPA is concurrently increased, and that the outcome is reflective of this overall cumulative change in enzyme activity. Potential candidates could include the following, which were all identified as altered in the HEK293 dataset: *PLA2G7, LPCAT3, LPCAT1* (all generate lysoPL), *PLPP1, PLPP2, PLPPR2* (all metabolise lysoPA).

Using the HEK293 dataset, we identified several enzymes that metabolize these lipids, and are known to be expressed in leukocytes, platelets, erythrocytes, heart, adipose tissue and plasma. We first focused on ATX (*ENPP2*), a plasma enzyme that converts primarily unsaturated lysoPC to lysoPA in healthy subjects (Table 4) ^46^. However, ATX was lower (Figure 2 C), and lysoPC metabolism by ATX appeared suppressed in the GG group (Figure 3 A-K). Furthermore, all lysoPCs were impacted by GG, including both saturated and unsaturated. Conversely, *ENPP2* was elevated *in vitro* in HEK cells. However, the HEK dataset only measures the impact of *ANRIL* knockdown on basal expression of *ENPP2* and it has been reported that inflammatory cytokine induction of *ENPP2* is suppressed by 50% in primary human monocyte derived macrophages that carry the 9p21 risk haplotype allele^47^. The role of ATX in CHD is not understood, and may vary with underlying genetic cause^48^. Indeed, while it metabolizes lysoPL to lysoPA in health, in acute coronary syndromes other pathways appear to predominate ^46^. Here, we found significant downregulation of ATX in GG plasma from middle-aged men who are otherwise healthy and without clinically detectable CHD (Figure 2 C). Thus, switching to ATX-independent metabolism may precede cardiovascular events in this risk group.

In healthy subjects, lysoPLs, particularly lysoPC, circulate at relatively high concentrations, where they could be generated by (i) lipases bound to the cell surface of endothelial cells in liver, heart and adipose tissues (*LPL, LIPC, LIPG*), (ii) Land’s cycle enzymes in circulating blood cells/platelets ^25^, (iii) lecithin-cholesterol acyl transferase (*LCAT*) trans-esterification in the liver, or (iv) by remodelling pathways for platelet activating factor (PAF) removal (Scheme 1). In healthy tissue, lipases predominate, but during vascular inflammation the balance may alter. The Land’s cycle involves phospholipase A2 (PLA2) hydrolysis, although the isoforms controlling blood levels are unknown. Candidates include stromal isoforms and cellular or secreted PLA2s from circulating cells and platelets. Also, a role for circulating/platelet PLA1 from platelets in lysoPL formation has also been proposed ^49^. In HEK cells, downregulation of *PNPLA2* (*ATGL*, a lipase) and *PLA2G4C* (PLA2Group IVC, a PLA2 strongly expressed in artery and heart) is consistent with our lipidomics findings of decreased lysoPL (Table 4) ^50, 51^ (Supplementary Data.xlsx, tab 7, Table 4, Scheme 1).

We also tested for upregulation of potential lysoPL removal pathways following *ANRIL* knockdown. LysoPL can be metabolized by *PLBD1* (Phospholipase B Domain Containing 1, expressed in neutrophils), removing the phospholipid headgroup, and this was significantly elevated in GG (Table 4, Scheme 1) ^52^. LysoPL can also be recycled back into PL pools via Land’s cycle enzymes, and consistent with this, upregulated *LPCAT2* (PC-acyl transferase in blood cells) and *MBOAT2* (a PE-acyl transferase in neutrophils) were seen (Table 4, Scheme 1) ^50, 53, 54^. Also, significant upregulation of *ACSL6*, a long chain acyl-CoA synthetase expressed in leukocytes and erythrocytes, required for fatty acid re-acylation was noted (Table 4, Scheme 1) ^50, 55-57^. *ACSL6* works in concert with *LPCAT2* and *MBOAT2*.

GG plasma showed significantly lower lysoPAs (Figure 2 D). These can be removed by phospholipid phosphatases, including *PLPPR2, PLPP1* (expressed in platelets), or *PLPP2* (in lung), and all were significantly induced by ANRIL knockdown ^58, 59^. In summary, our *in vitro* analysis provided several new candidates for reducing lysoPL/lysoPA in the context of Chr9p21-mediated CHD risk, including *MBOAT2, LPCAT2, ACSL6, PNPLA2, PLA2G4C, PLBD1, PLPP1, PLPP2* and *PLPPR2*. Based on their known cellular localization, potential sources are proposed (Figure 5 C)

LysoPLs have well characterized *in vitro* bioactivities, through mediating G protein-coupled receptor (GPCR) signalling that causes immune cell migration and apoptosis. This has led them to be proposed as “pro-inflammatory” ^49, 60-63^. However, most lysoPL is bound to albumin, immunoglobulins and other plasma carrier systems, and levels are already higher than required for mediating GPCR activation ^64, 65^. Lower levels of lysoPL in GG plasma suggests they are not pro-atherogenic in this case.

In addition to lipid class-specific changes in phospholipids, many significantly-decreased “unknowns” were found, which are currently absent in databases (Figure 1). The plasma lipidome contains large numbers of such species and a significant challenge lies in their structural and biological characterization. The comprehensive list of all lipids detected with fold-change and significance levels is provided (Supplementary data.xlsx, tab 1) as a resource for further mining.

A final question relates to how *ANRIL* and lysoPLs are functionally connected. PL metabolism is finely tuned during cell proliferation, with higher concentrations of lysoPC and lysoPE detected at G2/M, that then fall dramatically along with concomitant increases in PC/PE due to acylation during progression to G1 ^66^. This provides the PL membranes required to complete the cell cycle. Given *ANRIL’s* ability to regulate cell proliferation, and observations that silencing *ANRIL* prevents division and promotes senescence, the lower levels of lysoPLs in plasma may simply reflect altered rates of cell turnover in the vasculature, but this remains to be determined (Figure 5 C,D).

In summary, we reveal a selective association of altered GPL metabolism with CHD risk in a common risk SNP. In support of our findings, lower lysoPCs are associated with CHD factors such as visceral obesity, and a trend towards higher future risk of acute coronary events, although since lysoPA cannot be measured using shotgun or untargeted methods, we have not found other cohort data that includes this lipid as yet ^33, 37, 38^. In the Bruneck cohort, inclusion of lysoPCs in classifiers improved power for CHD risk prediction, indicating that although the reduction is rather modest, it is clinically significant^37^. Furthermore, the Malmö cohort reported that CVD development is preceded by reduced levels of lysoPCs^67^. Like our study, the reductions were rather modest being only around 8% for each lipid. In Malmö and Bruneck, the lipidomics methods were not as highly validated as in this study, thus our new data provides stronger analytical confidence while linking their findings to a specific risk locus and. Specifically, Bruneck applied a shotgun method with precursor/neutral loss scanning, while Malmö used untargeted MS and did not undertake any validation. Given the prevalence of rs10757274 GG in the general population (∼23%), our data may at least in part explain the findings in Bruneck and Malmö, with lower lysoPLs associating with a sub-group with a common SNP. Last, the alterations in multiple GPL regulatory pathways seen on *ANRIL* silencing, or the presence/removal of the risk locus *in vitro*, further indicate the involvement of bioactive lipids in this form of vascular disease, and mechanistic studies are warranted. To this end, fresh blood from AA and GG subjects is required to measure plasma and cellular levels of all candidate enzymes and relevant lipids, in order to identify how lysoPC/lysoPA metabolism is altered by the presence of the risk SNP. It would also be interesting to compare AG with AA and GG men, although for this, considerably larger sample numbers would be required than we have available currently in NPHSII.

## Supporting information

Supplementary Data.xls

## Author contributions

SM, JH, DW, PR, AE, MA, YZ, AC, CB, JAJ, RA, VT, CH, DS, JAoki, KK conducted experiments and undertook data analysis. SA supervised computational tool design. JAcharia, JC and JM collected and processed clinical samples. SM, VOD and SH conceived the experiments, designed the studies, and drafted the manuscript. All authors edited and approved the manuscript.

## Acknowledgements

We gratefully acknowledge expert discussion with Gerhard Liebisch (Regensberg), during revision of the manuscript.

## Figure Legends

**Scheme 1.**
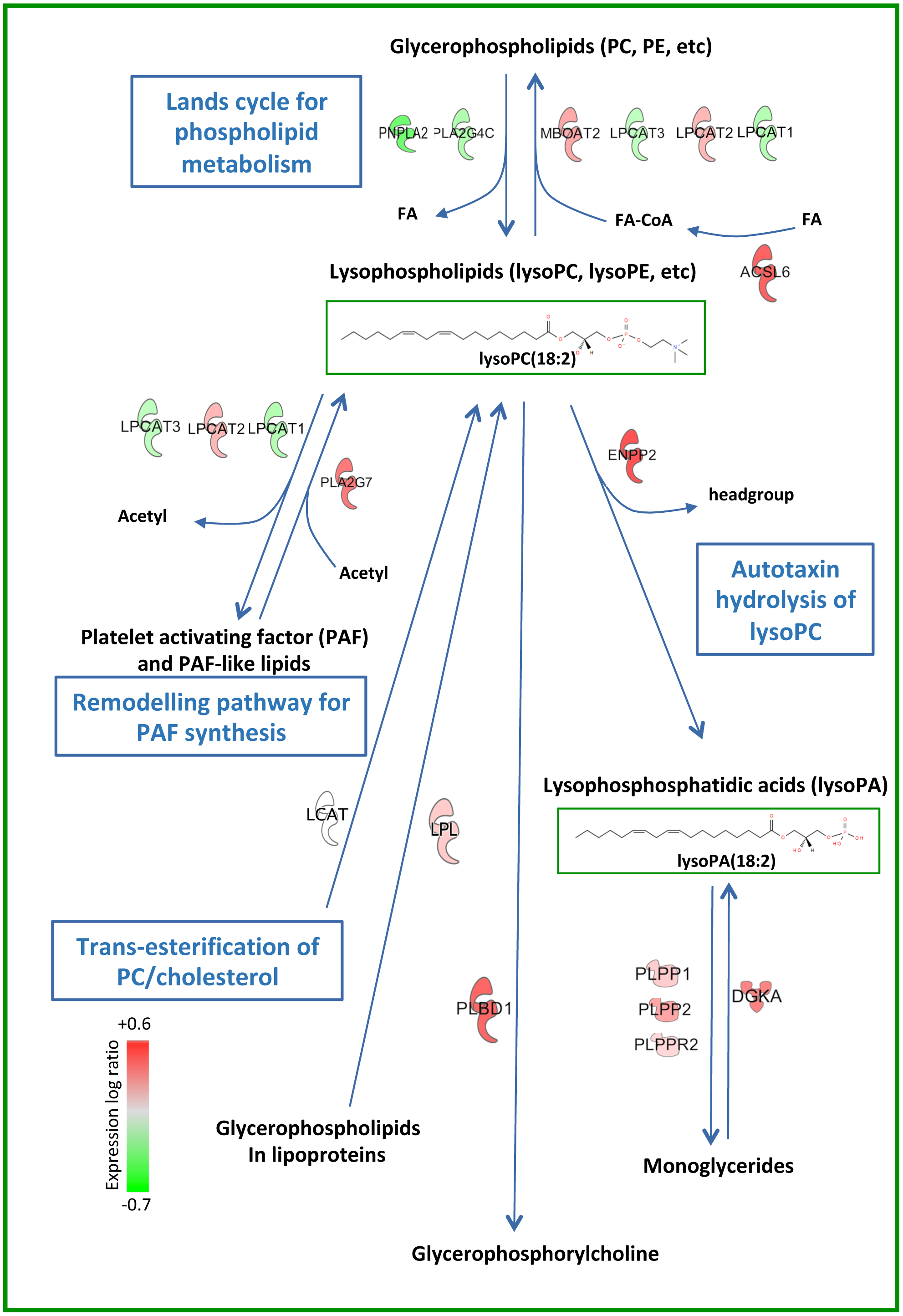
Metabolic pathway showing lysoPL/lysoPA regulatory genes that are significantly-altered by ANRIL knockdown. Genes that metabolise these lipids are shown. Full data on their transcriptional regulation is provided in Table 4. LCAT was not significantly regulated, but is shown for completeness.

