## Supplementary Data for "Metabolic dysregulation of the lysophospholipid/autotaxin axis in the chromosome 9p21 gene SNP rs10757274"

### Supplementary Methods and Data

### Supplementary Methods

*Patient samples*

NPHSII is a prospective CHD study of ~3000 men (20,21). Briefly, middle-aged men (aged 50–64 yrs) were recruited from 9 general practices in the UK 27 yrs ago. Exclusions included a history of CHD or diabetes. CHD was defined as acute myocardial infarction (MI), silent MI or undergoing coronary surgery. Ethical approval was provided by the National Hospital for Neurology and Neurosurgery and the Institute of Neurology Joint Research Ethics Committee, and Joint UCL/UCLH Committee of Human Research, Committees A and Alpha, and all samples were obtained with informed consent. There was a median of 13.5 years follow-up. Plasma lipids were measured at recruitment using standard methods as described[1]. Samples were selected which had been previously genotyped and shown to be carriers or homozygous for a particular gene variant (Table 1) [1, 2]. All plasma used had been stored at -80oC since isolation. SNPs were chosen based on known association with altered risk of CHD. For the *APOA5* and *CDKN2A/2B* SNPs, the rare allele is associated with elevated risk of CHD. The rare allele of the *SORT1* and *LDLR* SNPs are associated with lower levels of LDL-C and lower CHD risk. Carriage of the *APOE* E2 allele is associated with decreased risk, and the *APOE* E4 allele with higher risk of both CHD and Alzheimer's disease. Samples selected as controls were non-carriers for all the other selected risk gene variants, and all were homozygous for the *APOE* E3 allele. Thus, they were an appropriate comparison group for the other *APOE* genotype groups and the other SNP genotypes. Details of sample sizes, genes, SNPs and average levels of total triacylglycerides (TAG) and total cholesterol determined for these samples, are provided in Table 1.

*Global Lipidomics.*

To deep mine the human plasma lipidome, aiming to identify lipids that stratify with tissue type, disease state or treatment (including new species), an in-depth MS analysis is required. The full workflow for global lipidomics was performed in several separate "experiments" to enable regular calibration. Each consisted of samples with a known risk SNP, controls randomly chosen from the cohort, quality controls and blanks. Pooled control plasma was prepared using blood from ten healthy subjects. Samples were randomized before extraction. Control pooled plasma, water blanks and methanol blanks were analysed in a group after every tenth cohort sample. After lipid extraction the samples were analysed in the same randomized order as the extraction using LC-ESI- FTMS in positive and negative mode. A 50 min separation in high-resolution mode was utilised. All major lipid classes, including low abundance species such as fatty acids (FA)/eicosanoids were detected. These global lipidomic datasets are of significant size requiring specialised informatics. Extensive clean up is used to remove artefacts (adducts, background contaminants, etc), combining positive and negative runs to generate single datasets for comparison between sample sets, and putative identification and assignment to specific lipid classes. This presents an enormous informatics challenge, for which we recently generated the Python workflow LipidFinder[3]. Lipids were extracted by combining 950 µL water with 50 µL plasma, 4 µl glacial acetic acid and 250 pg internal standards (2H8-arachidonic acid (Cayman Chemicals) and 4ME 16:0 Diether PE) (Avanti Polar Lipids)). Next, 2.5 mL of solvent (1M acetic acid/propan-2-ol/hexane; 2:20:30; v/v/v) was added and samples vortexed for 1 min. Hexane (2.5 mL) was added, vortexed for 1 min and centrifuged for 5 min at 500 g, 4°C. The upper hexane layer was collected. The sample was re-extracted by adding hexane (2.5 mL) to the remaining aqueous phase, vortexing for 1 min and centrifuging for 5 min at 500 g, 4°C. Again, the upper hexane layer was collected and combined with the first hexane layer. The remaining aqueous phase was re-extracted using a modified Bligh and Dyer protocol[4]. 3.75 mL solvent (chloroform/methanol; 1:2; v/v) was added per sample. After vortexing for 1 min., 1.25 mL chloroform was added, and vortexed again for 30 sec., before adding 1.25 mL water, followed by 30 sec. vortex. Samples were centrifuged for 5 min. at 500 g and 4°C and the bottom chloroform layer collected and combined with the two hexane layers before drying under vacuum. Samples were re- suspended in 400 µL methanol and filtered through a centrifuge filter (10 kDa cut off, Millipore) before LC-MS.

Global lipidomics was carried out on an Accela liquid chromatography system coupled to an Orbitrap Elite mass spectrometer (Thermo Fisher Scientific). Liquid chromatographic separation was performed at 30°C using an Accucore C18 (Thermo Fisher Scientific) reversed phase column (150 × 2.1 mm, 2.7 µm) with solvent gradient of mobile phase A (water/acetonitrile; 80/20; v/v; 4 mM ammonium acetate and 0.1 % glacial acetic acid) and B (acetonitrile/isopropanol; 30/70; v/v; 4 mM ammonium acetate and 0.1 % glacial acetic acid) at 0.4 mL/min over 60 min. The linear gradient of B was 16 - 60% for 12 min, 60 - 72% from 12 min - 19 min, 72 - 84% from 19 min - 42 min, 84-98% from 42 min - 51 min and held at 98% for another 6.5 min before re-equilibration. MS conditions were as follows for analysis in positive ESI ionization mode: resolution 60,000 at 400 *m/z* (providing approx. 3 scans/sec) HESI-II temperature 400°C, N2 as drying gas, sheath gas flow 37 arbitrary units (au), auxiliary gas flow 15 au, sweep gas flow 1 au, capillary temperature 320°C, spray voltage + 4.0 kV, S-lens RF level 62 %. Lock mass was *m/z* 391.2843. For analysis in negative ESI ionization mode: resolution 60,000 at 400 *m/z*, HESI-II temperature 350°C, N2 as drying gas, sheath gas flow 37 au, auxiliary gas flow 15 au, sweep gas flow 2 au, capillary temperature 320 °C, spray voltage – 3.5 kV, S-lens RF level 69 %. Lock mass was *m/z* 265.1479.

*Informatics and statistical analysis of global datasets.*
Orbitrap datasets were processed using the R version of XCMS (Version 3.4) for feature detection and alignment[5]. Extracted and aligned features were further processed using LipidFinder to remove isotopes, adducts and background contaminations, with parameters as in Supplementary Methods[3]. Missing values were set at zero. In addition to using Websearch in LipidFinder, with only curated lipids from LIPID MAPS, we applied an in-house generated retention time and m/z database which identifies around 1,000 lipids known to be in human plasma (level three identification according to the metabolomics standard initiative[6]). The LIPID MAPS database enabled a lipid category to be assigned, with a putative match generated where possible. Settings for the programs used and database lipids are provided as a supplementary file (databasemethod.xls). This enabled assignment of a putative lipid class to 30 – 50 % of all ions detected. Next, univariate statistics were applied to identify interesting lipid features (Mann-Whitney u test (non-parametric), calculation of fold change). We chose Mann Whitney U because we expect that some features will deviate from normality. Chromatograms for all lipids with $p \leq 0.075$ or a high relative fold-change (top 20 % of the features) were manually checked to ensure peak quality, by examining raw data Xcalibur datasets (around 1,500). Next, we processed the

data using quantile normalization, applied non-parametric Mann–Whitney U test assuming unequal variance and adjusted the resulting p-values to confirm significance. Quantile normalization was performed using the normalize.quantiles function in the preprocessCore package in R (Version 1.4)[7]. p-Values were adjusted for lipid classes by the sequential goodness of fit approach described in by using the R based version (Version 2.3) [8].

*Targeted analysis of TGs, CEs and free cholesterol.*
Plasma was extracted as for global lipidomics, outlined in Supplementary Methods. LC/MS/MS was performed on a Nexera liquid chromatography system (Shimadzu) coupled to an API 4000 qTrap mass spectrometer (Sciex). Plasma (5 µL) was added to 500 µl water, and 10 µL internal standards (50 ng 2H5-TG(51:1), 500 ng 2H7-CE(18-1)) added. 1.25 mL of solvent (1M acetic acid/propan-2-ol/hexane; 2:20:30; v/v/v) was added and the sample vortexed for 1 min. Hexane (1.25 mL) was added, vortexed for 1 min and centrifuged for 5 min at 500 g, 4°C. The upper hexane layer was collected. The sample was re-extracted using hexane, vortexed for 1 min and centrifuged for 5 min at 500 g, 4°C. Again, the upper hexane layer was collected and combined with the first. The aqueous phase was then re-extracted using a modified Bligh and Dyer protocol[4]. 1.9 mL solvent (chloroform/methanol; 1:2; v/v) was added. After vortexing for 1 min., 0.6 mL of chloroform was added, and vortexed again for 30 sec. before adding 0.6 mL water, followed by 30 sec. vortexing. Samples were centrifuged for 5 min. at 500 g, 4°C and the bottom chloroform layer collected and combined with the two hexane layers before drying under vacuum, and re-suspending in 400 µL methanol and storing at -80 °C until analysis. Lipid extracts were diluted 1:10 with MeOH. LC-MS/MS for free cholesterol and cholesterol esters and LC-MS analysis of triacylglycerides was performed on a Nexera liquid chromatography system (Shimadzu) coupled to an API 4000 qTrap mass spectrometer (Sciex). Liquid chromatography was performed at 40 °C using a Hypersil Gold C18 (Thermo Fisher Scientific) reversed phase column (100 × 2.1 mm, 1.9 µm) at a flow rate of 0.4 mL/min over 11 min. Mobile phase A was (water/solvent B 95/5; v/v and 4 mM ammonium acetate) and mobile phase B was acetonitrile/isopropanol (40/60; v/v and 4 mM ammonium acetate). The following linear gradient for B was applied: 90 % for 1 min, 90 – 100 % from 1 to 5 min and held at 100 % for 3 min followed by 3 min at initial condition for column re-equilibration. Triglycerides were analysed in selected ion monitoring (SIM) mode covering a range from TG(32:0) up to TG(56:0) including also unsaturated TGs (Supplementary Table 1). MS conditions were: ESI temperature 450°C, N2 as drying gas, ion source gas1 35 psi, ion source gas2 50 psi, curtain gas 35 psi, ESI positive spray voltage 5,0 kV, declustering potential 60 V and entrance potential 10 V. Dwell time was 10 ms resulting in a cycle time of 0.56 sec. TAGs were quantified using an external calibration with the following TG species (TG(14:0-16:1-14:0)-d5, TG(15:0- 18:1-15:0)-d5, TG(16:0-18:0-16:0)-d5, TG(19:0-12:0-19:0)-d5 and TG(17:0-17:1-17:0)- d5). Free cholesterol and CEs were analysed in MRM mode monitoring the parent to daughter transitions of 12 CEs and free cholesterol (Supplementary Table 2). MS conditions were as follows: ESI temperature 150°C, N2 as drying gas, ion source gas1 25 psi, ion source gas2 50 psi, curtain gas 35 psi, ESI positive spray voltage 5.0 kV, declustering potential 70 V, entrance potential 10 V, collision energy 20 V, and collision cell exit potential 25 V. Dwell time was 75 ms for each transition and the cycle time 1.12 sec. cholesterol and CEs were quantified using an external calibration with the following CE standards: CE(14:0), CE(16:0), CE(18:0), CE(18:1), CE(20:4), CE(22:6) and CE(18:1-d7).

*Targeted analysis of lysoPLs .*

Plasma (50 µL) was added to 950 µl water, and 10 µL internal standards (20 ng LysoPC(18:1-d7) and LysoPE(18:1-d7) and 4µL glacial acetic acid added. 2.5mL of solvent (1M acetic acid/propan-2-ol/hexane; 2:20:30; v/v/v) was added and the sample vortexed for 1 min. Hexane (2.5 mL) was added, vortexed for 1 min and centrifuged for 5 min at 500 g, 4°C. The upper hexane layer was collected. The sample was re-extracted using hexane, vortexed for 1 min and centrifuged for 5 min at 500 g, 4°C. Again, the upper hexane layer was collected and combined with the first. The aqueous phase was then re-extracted using a modified Bligh and Dyer protocol[2]. 3.75 mL solvent (chloroform/methanol; 1:2; v/v) was added. After vortexing for 1 min., 1.25 mL of chloroform was added, and vortexed again for 30 sec. before adding 1.25 mL water, followed by 30 sec. vortexing. Samples were centrifuged for 5 min. at 500 g, 4°C and the bottom chloroform layer collected and combined with the two hexane layers before drying under vacuum, and re-suspending in 400 µL methanol. Samples were filtered (Amicon® Ultra centrifugal filter units, 10,000 NMWL; Millipore) before storing at -80 °C until analysis. LC-MS/MS was performed on a Nexera liquid chromatography system (Shimadzu) coupled to an API 4000 qTrap mass spectrometer (Sciex). Liquid chromatography was performed at 30 °C using a Accucore C30 (Thermo Fisher Scientific) reversed phase column (100 × 3.0 mm, 2.6 µm) at a flow rate of 0.5 mL/min over 52 min. Mobile phase A was acetonitrile/water (20/80 v/v; 5mM ammonium acetate; 0.1% v/v glacial acetic acid) and mobile phase B was acetonitrile/isopropanol (30/70 v/v; 5 mM ammonium acetate; 0.1% glacial acetic acid). The following linear gradient for B was applied: 60 % for 0.5 min, 60 – 90 % from 0.5 to 15.5 min and held at 90 % for 40 min followed by 10 min at initial conditions for column re-equilibration. LysoPL were analysed in MRM mode monitoring the parent to daughter transitions of 8 lysoPC and 7 lysoPE species (Supplementary Table 3). MS conditions were: ESI temperature 450°C, N2 as drying gas, ion source gas1 40 psi, ion source gas2 30 psi, curtain gas 20 psi, ESI negative spray voltage 4.5 kV. Dwell time was 100 ms resulting in a cycle time of 2.8 sec. Entrance potential was 10 V and collision energy 36 V. Declustering potential was 165 V and 112 V for lyso PC and lyso PE species, respectively. LysoPL were quantified using an external calibration using various species (lysoPC(16:0), lysoPC (18:0), lysoPE(16:0), lysoPE(18:1)) in a concentration range between 0.1-1000 ng/mL.

*Analysis of Affymetrix data from ANRIL down-regulation in cell lines*

Raw Affymetrix CEL files relating to total transcript expression in HEK 293 cells stimulated with Tetracycline (shRNA ANRIL silenced for 0h, 48h and 96h) were downloaded from the GEO database (accession: GSE111843) and analysed using packages in CRAN and Bioconductor: limma, oligo, ggplot2 [9-13]. Data were processed using RMA-normalization and differential gene expression analysis performed using “best practice” detailed in the limma vignette. P-values were corrected for multiple- testing using Benjamini-Hochberg. GO-term enrichment analyses for significant differentially expressed genes (adjusted p-value cut-off: 0.05) were performed using the online PANTHER enrichment tool[14]. The IPA software (QIAGEN Inc.) was used to generate gene/protein interaction networks from genes identified as being significantly differentially expressed (between 96h vs 0h, and 48h vs 0h) and mapping to a common lipid process-associated GO-term. Networks were plotted within IPA using the standard Ingenuity Knowledge Base and default analysis settings. Volcano plots were generated in R using ggplot2.

*RNAseq Methods and Analyses for iPSC-derived Vascular Smooth Muscle Cells* iPSCs were created from patient-derived peripheral blood monocytes, edited to selectively delete the locus, differentiated, and lysed, prepared, and run on an Illumina HiSeq2500 to obtain transcriptomic expression data from RNA-sequencing as reported by Lo Sardo and coworker[15]. Data was normalized using variance stabilizing transformation and differentially expressed genes (DEGs) were identified using DESeq2 R package (v1.8.2)[16]. Data was selectively clustered using lipid enzyme genes reported in Supplementary Data.xlsx, tab 7, by MORPHEUS[17], employing hierarchical cluster with Pearson correlation to identify DEGs with common expression patterns. These data were further analysed for principal components using ClustVis in R[18]. Data is grouped and reported by patient genotype for the non-risk haplotype (NNWT), risk haplotype (RRWT) and their genome edited counterparts (NNKO and RRKO). DEGs where RRWT is lower than the other three genotypes were identified and significance assessed by Mann Whitney U test.

**XCMS parameters and R-script**

```
# all samples

setwd("C:/myData")

myClass1 <- "c"

myClass2 <-"sol"


# peak picking using wavelet algorithm for peak detection (centWave)

xset <- xcmsSet (method="centWave",ppm=10, peakwidth=c(10,120), snthresh=5,
prefilter=c(10,20000), integrate=1, mzdiff=0.001, fitgauss=FALSE,  noise=20000,
scanrange=c(1,11485))


# peak alignment

xset <- group(xset, bw=30, mzwid=0.005, minfrac=0.5, minsamp=1)


# retention time correction

xset <- retcor(xset, method="obiwarp", profStep=0.05, response=20, center=1
,plottype="deviation")


#re-align

xset <- group(xset, bw=3, mzwid=0.005, minfrac=0.5, minsamp=1)


# fill in missing peak data

xset <- fillPeaks(xset)


# output results

reporttab <- diffreport(xset, filebase="output")
```

### LipidFinder Parameters

| Parameter | Parameter descritption | Expected Data Type | Current value |
|---|---|---|---|
| firstRepOffset | This is the index of the first sample, first replicate column | Integer(>3) | 3 |
| numberOfSamples | The number of samples in the experiment | integer (>0) | 49 |
| numberOfTechReps | The number of replicates of each sample | integer (>0) | 1 |
| numberOfQCReps | The number of QC replicates in the input file(s) | integer (>=0) | 0 |
| numberOfSolventReps | The number of solvent replicates in the input file(s) | integer (>=0) | 5 |
| filePolarityMode | File Polarity Mode (P: Positive mode files | | |
| columnType | The type of column used for LC (PO: Polar | | |
| QCLowRSD | Lower relative standard deviation cut off | integer (>0 AND <QCHighRSD) | 30 |
| QCHighRSD | upper relative standard deviation cut off | integer (>QCLowRSD AND <100 | 50 |
| removeSolvent | Solvent removal toggle (TRUE: Remove solvent intensity | | |
| solventFoldCutOff | The minimum fold difference greater than the solvent intensity a sample replicate intensity must be in order to be considered significant enough to process further | Float (>0.0) | 3 |
| intensitySignificanceCutOff | The level at which the intensity of a sample reading is significant enough to process | Integer (>0) | 1 |
| mzFixedError | The fixed error allowable when observing a mass | Float (>0.0) | 0.0005 |
| mzSizeErrorPPM | The mass size dependant PPM error to add to the fixed error | integer (>0) | 4 |
| peakMaxRTWidth | The maximum allowable retention time (mins) a single lipid peak can span | Float (>=3*peakAdjacentFrameMaxRT ) | 0.003 |

| Parameter | Parameter descritption | Expected Data Type | Current value |
|---|---|---|---|
| peakMinFoldCutOff | The minimum fold difference greater than the adjacent candidate frame's intensity that the current frame intensity must be in order for the current frame to be considered part of a peak. | Float (>1.0) | 1.3 |
| peakAdjacentFrameMaxRT | The maximum time difference (mins) between a feature edge and an adjacent frame where the adjacent frame could be considered for inclusion in the same feature | Float(<=peakMaxRTWidth/3) | 0.001 |
| peakConcatenateAllFrames | This toggle indicates which peak concatenation should be performed (TRUE:Concatenate all peak frame intensities into the peak centre, FALSE:Concatenate only the most intense peak frame into the peak centre) | boolean (True or False) | TRUE |
| removeContaminant | This toggle allows the user to specify whether contaminant removal should be executed (TRUE: Remove contaminants | | |
| removeAdduct | This toggle allows the user to specify whether adduct removal should be executed (TRUE: Remove adducts | | |
| adductAddition | Adduct addition toggle. Allows the user to specify whether they wish to add the intensity of any adducts identified to the intensity of the primary mass (True: Add adduct intensity to primary mass intensity | | |
| removeStack | This toggle allows the user to specify whether stack removal should be executed (TRUE: Remove stacks | | |
| maxStackGap | The maximum number missing values before a stack search will terminate for a particualr lipid or contaminant | integer (>=0) | 3 |
| lipidStackAddition | Lipid stack addition toggle. Allows the user to specify whether they wish to add the intensity of any lipid stacks ions identified to the intensity of the primary mass (True: Add adduct intensity to primary mass intensity | | |
| rtTolMultipler | A multiplier for peakAdjacentFrameMaxRT to allow a smaller tolerance in certain circumstances (e.g. when looking for stacks) | Float (<=1.0) | 1 |
| outlierHighIntensityValue | The cut off point of the replicate means of a sample between using the lower RSD cut off (outlierLowIntensityRSD) and the higher RSD cut off (outlierHighIntensityRSD) | integer (>0) | 5000 |
| outlierLowIntensityRSD | The RSD to use when the mean average intensities are lower than the replicate mean intensity cut off (repMeanOutCorCutOff) | integer (>0 AND <outlierHighIntensityRSD) | 35 |
| outlierHighIntensityRSD | The RSD to use when the mean average intensities are higher than the replicate mean intensity cut off (repMeanOutCorCutOff) | integer (>outlierLowIntensityRSD) | 40 |
| featureLevelMassAssignment | This toggle allows the user to specify whether they wish to assign the masses of every frame to the highest intesity mass in the feature set, default is assignment at the mass group level (TRUE: Re-assign every | | |

| Parameter | Parameter descritption | Expected Data Type | Current value |
|---|---|---|---|
| | mass in a feature group to the mass of the highest intensity frame within the feature group | | |
| negativeModeAdductPairs | The pairs of negative adducts relating to the index in the adducts.csv file | 2d list of indices example format [[0,1],[0,2],[3,4],_] | [[0,1],[0,2],[0,3],[0,4],[2,3]] |
| positiveModeAdductPairs | The pairs of positive adducts relating to the index in the adducts.csv file | 2d list of indices example format [[0,1],[0,2],[3,4],_] | [[5,6],[5,7],[5,8],[5,9],[5,10],[6,7],[6,10],[8,9]] |
| broadContsdMult | | | 1 |
| broadContminPoints | | | 4 |
| broadContRSDCutOff | | | 30 |
| broadContrtSDCutOff | | | 2 |
| retentionTimeLowCutOff | | | 1.3 |
| retentionTimeHighCutOff | | | 56 |
| rtCorrectStDev | | | 999 |
| rtCorrectMeans | | | FALSE |

### Supplementary Results

*Targeted MS analysis of triglycerides (TG), cholesterol and cholesteryl esters (CE) shows no change in profile between genotypes*

Traditional cardiovascular risk factors were measured many years ago for the NPHSII cohort. Overall, total cholesterol, TG and CE tended to be lower for this GG risk group (Table 1). This is a reflection of the control AA group randomly selected, which overall had basal lipid levels at the upper range of normal, since large cohort studies have shown that this risk variant is not associated with changes in circulating lipoproteins (1, 9). However, clinical measurements do not include the large number of molecular species of both TG and CE. Herein, quantitative MS analysis found that the relative abundance of these across genotypes is similar (Supplementary Figure 7). Lipidomics of TG generally reports single "species" based on molecular weight, which describes the total number of carbons and double bonds in the FAs, e.g. TG(52:4)[19]. Thus, we also interrogated individual TG peaks eluting within each chromatogram for genotype-specific changes, but none were found (data not shown). Thus, the profile of individual CE and TG molecular species are not altered in the GG risk genotype.

*Global lipidomics analysis of plasma*

The LipidFinder approach provides an extensive but unvalidated dataset, considered analogous to a gene array (Figure 1 A). This required extensive method development for plasma samples, outlined in full below.

(i) Optimization of the workflow. XCMS was used for initial processing, however this is not designed for deep mining to detect "unknown" lipids, and was unable to remove many artefacts. Post-XCMS, around 14,000 ions are retained in the dataset, far more than would be considered representative of true lipids (Figure 1 A). When LipidFinder was then used for automatic data clean-up and removal of duplicate ions in both positive and negative ion modes, the number reduced to around 10,000. This was followed by isotope removal, then a manual clean up including intensity cut-off and missing value analysis. Features with > 50 % missing values equally distributed across all samples were considered as artefacts/below LOD and removed. Next, extracted ion chromatograms (EIC) were generated for all ions with p-values ≤ 0.075 and/or high fold-change (top 10 % of features with highest change in either positive or negative mode) and corrected for noisy/abnormal peak shape/spikes. Any peaks not matching the quality criteria (> 3-times background level, Gaussian peak shape and <10 data points per peak) were removed. These steps further reduced the number of ions to around 2,000 - 3,000 per sample. Scatter diagrams show the difference between data obtained post-XCMS, then after LipidFinder and manual clean- up steps, including WebSearch and in-house database search (Figure 1 A,B). Major lipid categories were assigned according to the LIPID MAPS classification system. As expected for reverse phase chromatography, glycerophospholipids (GPL, green) elute before more non-polar glycerolipids (GL, including TGs, DGs, red) and sterols (including CEs, pink). The most abundant lipids detected were glycerophospholipids (GPLs, including phospholipids (PL) and lysoPL (lysoPL)) accounting for ~21%, followed by glycerolipids (GL, including diglycerides (DG) and triacylglyceridesTG, 12%), sphingolipids (SL, 9%), fatty acyls (FA, ~3%), sterol lipids (~1.4%) and prenol lipids (<1%). Unknowns accounted for 53%. An example sample analysis is shown in Figure 1 B.

(ii) Assay performance
Throughout the experiment total ion current (TIC) chromatograms for QC samples were visually compared, since these originate from a single pooled plasma sample. Also, extracted ion chromatograms (EIC) of several known lipids were compared for chromatographic peak shape and mass accuracy across all QC samples. % CV was calculated for all features and those with % CV higher than 50% were removed from the dataset. For example, average % CV across all features for the QC samples of the *CDKN2A/2B* dataset was 17.5 % (n = 7). In addition, % CV was calculated for 15 lipid classes (one lipid per class). As an example, Figure shows boxplots of these 15 lipids for the QC samples (Supplementary Figure 8). Here, % CV ranged from 5.7 % for γ-linolenic acid up to 28.9 % for PS(38:1), comparable with previous studies[6].

(iii) NPHSII plasmas are broadly similar in composition to fresh samples for most lipid classes. NPHSII plasma was collected several decades ago. Many cohorts archive serum, a biological fluid generated by clotting whole blood *in vitro*, during which white cells and platelets are activated generating a multitude of inflammatory lipids. Thus, only plasma was analysed herein. Additionally, incorrect storage temperatures may allow artefactual oxidation of unsaturated species. A random set of NPHSII controls was compared with plasma from genetically-unrelated healthy donors obtained in the present day (n = 10 for both) using untargeted MS. Total ion current (TIC) for NPHSII versus fresh samples was found to be similar (Supplementary Figure 1 A). Relative abundance of selected lipids from different classes was then compared[20]. The majority were not significantly different and likely represent normal variation in unrelated human subjects (Supplementary Figure 1 B). Also, several TGs and CEs were separately compared using a targeted assay, and only 1 of 17 was significantly lower in NPHSII samples (Supplementary Figure 2 A,B). Since oxidation during storage could lead to artefactual generation of bioactive species, we measured four non-enzymatically-oxidized PLs using targeted MS and found these to be considerably higher in the older samples (Supplementary Figure 2 C). Last, we measured lysoPC signals from the untargeted dataset and found that there were some increases, although not all were significant (Supplementary Figure 3). Thus, for most lipid classes, plasma that has been correctly stored for many years is usable. In the case of oxidized lipids and lysoPLs the impact of changes during storage need to be considered, including whether differences in lysoPLs could result from altered levels of plasma lipase enzymes inherent in plasmas from different patient groups (this is discussed in more detail below).

**Supplementary discussion on cohort lipidomics**

*Choice and interpretation of lipidomics based on methods used.*
Here, we used an untargeted analysis to generate a hypothesis which we next addressed using fully-validated gold standard quantitative targeted methods as a second step. We then followed up further, measuring again the lipids of interest in independent sets of samples. Generally, cohort lipidomic studies that measure large numbers of samples using untargeted methods do not take this rigorous approach. Instead they often report on lipids that have been only partly identified (e.g. a “putative” structure” only) and provide fold change values only, with no information on the amounts present in the samples. Thus they take a first step, analogous to a gene array, but many do not validate using appropriate targeted methods. While gaining popularity, this approach is unhelpful for lipidomics overall because it is leading to errors in reporting lipids and their stratification with disease risk,

phenotype or genotype. Also, actual values/amounts of lipids are not measured, meaning the results are considerably less informative and impossible to directly compare across studies.

It is essential to consider the untargeted approach in the right context in relation to our study. As background, there are several different lipidomic approaches available, and all have their pros and cons in terms of coverage of lipids, sensitivity and selectivity. Methods that are applied to large cohorts generally are (i) fast, due to short or no chromatography, and (ii) utilise high resolution MS (untargeted global lipidomics). However, there are major pitfalls with this approach:

First, short methods suffer from ion suppression, where low abundance molecules are not detected at all. Thus, one can only survey highly abundant lipids that comprise the circulating lipoprotein compartment (e.g. triglycerides, cholesterol esters). For lower level and often far more biologically important signalling lipids, e.g. those that would be involved in inflammation, long separations are required. However, increasing time of analysis per sample introduces the necessary trade-off of a method becoming low throughput.

A second issue relates to the limitations of high resolution MS. This approach is not quantitative and does not fully identify the lipid structures being measured. Untargeted MS only provides a "putative" structure (a best guess), and for some lipids there can be up to 30 or more molecular species in databases that could account for a single detected peak. Incorrectly annotating lipids using MS data is a major problem in the literature currently with many investigators new to the field sometimes "identifying" and reporting on lipids incorrectly using this type of approach. The specific problem relates to the misuse of databases with over-interpretation of untargeted data. For example, we see many studies where databases are being used to fully annotate lipid structures (e.g. down to stereochemical structures) despite only the molecular mass being obtained from an untargeted analysis. This is not best practice, and is currently a significant problem in the field leading to erroneous reporting of findings. We are currently addressing this for the benefit of the lipidomics/metabolomics community in LIPID MAPS (www.lipidmaps.org) and Lipidomics Standards Initiative (LSI) through developing a hierarchical database and new guidelines on the application and use of shorthand nomenclature for lipids (it is a lipid specific issue that does not impact non-lipid metabolites). Because of this issue, in our study herein, we did not annotate our untargeted MS data using lipid names. Instead we assigned ions to lipid categories, and then statistically analysed the categories separately to examined for SNP-dependent differences. We prefer this approach since it avoids over interpretation and rather than using untargeted to infer findings, we use that approach as an initial screen only.

Here, for the deep phenotyping of lipids including lower abundance and some signalling lipids, a tailored approach was used that first combined a long chromatography analysis with an untargeted global MS method on an Orbitrap platform, as our initial screen. Our long chromatography provides the best separation of individual molecular species, and will reveal changes that are simply not possible to detect using faster more high throughput methods, often used for large cohort analysis. Our idea was that we could detect low abundance lipids and also unknowns, and then develop hypotheses for validation. As this was a time consuming method, we analysed a relatively small number, then validated both in that sample set and a second set independently, using gold standard targeted methods. This approach is specifically designed to avoid such pitfalls as misidentification and false positives in studies involving smaller numbers, as well as to provide quantitative values for properly identified lipids. This approach is not feasible for large cohorts, but it is far more rigorous than screening large numbers and not validating at all, which is becoming common in the field. This study is thus designed to uncover pathways and networks as a first step, and then to stimulate further mechanistic study on these lipids in this context in humans. In summary, the untargeted method is being used as an initial screen only, to direct our fully validated explorations down to sub-classes of signalling phospholipids and their potential role in this form of CVD.

It is important to point out that untargeted methods cannot detect lysoPAs at all, and not all lysoPCs are detected either, due to lower sensitivity compared with targeted methods. Generating accurate data on these lipid classes requires specialised high sensitivity targeted assays such as used in this study. Also, ATX is measured using an antibody-based approach, not lipidomics.

As part of preparing this manuscript, we looked into the literature in detail and discussed our findings with a number of experts who work in this area, including Gerhard Liebisch who conducted the lipidomics analysis in[21] and Karsten Surhe who hosts a website collating known GWAS/metabolomics studies. Unfortunately, we have not able to gather further information relating to what lipids might be altered in other cohorts with this SNP. Either the patient demographics or the diseases being studied were different, lysoPCs or lysoPAs were not measured, or there were concerns relating to artefactual increases in lysoPL due to sample processing considerations (see later for more details). Where untargeted data was available, no quantitative information on the lipids was available thus generation of lysoPLs during storage was impossible to assess.

In relation to our mechanistic studies in plasma, we found evidence that the metabolism of lysoPL and lysoPA is altered in AA versus GG samples. To take this finding further, the next step would need to be a new cohort study to determine the specific enzymes (from those we have identified in the HEK293 or VSMC datasets) that might be responsible for plasma lysoPL and lysoPA levels in humans, and how they alter with this particular SNP. We note that the cellular origin of lysoPL and lysoPA in plasma is not well understood, and that resolving this is an important question that should follow our study. An approach would be obtain fresh whole blood and profile levels of all candidate enzymes in circulating cells. This is a major project requiring funding and would take a number of years (needing ethical approval, cohort collection, genotyping, recruitment, sampling, analysis, etc). Determining vascular tissue or liver levels of enzymes in humans would also be relevant, but at this time isn't feasible. Our dataset provides a first step towards this that we expect to be followed up by us, and we hope also by other investigators.

Our initial untargeted experiment was relatively small because we preferred to undertake a deep phenotyping study as a first hypothesis-generating step, using long chromatography analysis, which by its nature is low-throughput. To then address this issue, we used a fully validated targeted assay, which analysed lysoPL alone, and in that case we showed that all lysoPCs measured were statistically significantly lower. LysoPLs were in general normally distributed (Kolmogorov-Smirnov test) or near normal, while ATX was normal. For both lysoPLs and ATX, we include both t-tests (black) and Mann Whitney U (red) on our box and whisker plots, and note that we get virtually identical results with either test (in some cases, the Mann Whitney U test returns higher levels of significance for AA versus GG).

To further examine the issue of statistical power, we conducted a power calculation for all variables using GPower[22]. These were performed for a t-test with alpha = 0.05 and power at 80%. Minimum sample numbers returned varied from 38-71 for lysoPCs (e.g. 16:0 - 44, 18:0 - 38, 18:1 - 44, 18:2 - 71 for lysoPCs), and 23 for ATX. Generally, non-parametric tests would require up to 15% more, so the sample sizes for lysoPCs and ATX were sufficient.

For lysoPA, we first analysed the second validation cohort (just under 50 samples per group). However, our power calculation indicated numbers were required as follows: 14:0 – 65, 16:0 – 52, 16:1 – 53, 18:0 – 81, 18:1 – 69, 18:2 – 60, 18:3 – 201, 20:3 – 36, 20:4 – 39, 20:5 – 110, 22:6 – 3000). Furthermore, lysoPAs were not normally distributed since they had some values that tended to be considered “outliers” on the higher side of the median. This was more obvious for AA than GG samples in general. To address this, we analysed a second cohort of samples from different individuals, bringing the total number of samples to 95,100 (AA,GG), and providing sufficient statistical power for 8 out of 11 lysoPAs. It is well accepted that t-tests can be applied to non-normal data, so long as the sample size is sufficiently large. There is debate in the literature on what sample size would be considered sufficient with estimates from 15 up to 80 in various papers [23]. As we have 95-100 for our groups, we have applied both t-test and Mann Whitney U test. We found that using t-test, that 7 of 11 lysoPAs are significantly lower in the GG risk group, while we did not see significant differences using Mann Whitney U. Thus, overall lysoPAs were lower, but extent of the reduction was relatively small.

*Issues of sample storage and processing relating to lipid levels.*
We took significant efforts to ensure that the cohort plasma was of sufficient quality. This is not commonly done in the literature, but it is critically important for cohort studies, and should be strongly encouraged. In comparison with fresh plasma, we found that most lipids were unchanged, however oxidized phospholipids and lysoPCs were somewhat increased during the time in storage (lysoPA is not detectable using untargeted lipidomics). This is not perhaps totally surprising because it is known that lysoPCs in plasma are influenced by sample collection, anti-coagulation and time to freezing, and oxidation over a long period is to be expected to some degree at least. Relating to lysoPCs, if plasma is kept at room temperature for a day or more, these lipids increase significantly[24]. In the case of NPHSII samples used herein, all samples were centrifuged at 1000 x G for 10 min at room temperature before immediate low temperature storage. This will have prevented their generation during initial sample processing. However, it is likely that during the 25 yrs of storage, some generation of these lipids will have occurred at a low rate. Although we could not measure this in the Orbitrap data, similar elevations in lysoPA may have occurred, as Liebisch has also seen that these lipids also rise with storage of plasma at room temperature[25]. The enzymes responsible for this are not known, but ATX is a likely contributor, along with plasma PAF-AH (PLA2G7) which generates lysoPC from oxPL (lipids which we also found were increased in storage).

Since lysoPC appears to have increased in storage, we cannot say for certain that the altered levels we found in AA vs GG were already present in vivo, or if this occurred during low temperature storage. However, regardless of this, the AA and GG plasma is different in relation to both lysoPC and lysoPA and their underlying metabolism, and elucidating why is important in terms of revealing the underlying vascular lipidomic impact of this SNP. Our observations need to be considered a starting point for further studies to fully delineate the biological processes involved. We also point out that our findings are relevant for others undertaking cohort studies some of which are uncovering similar alterations in lysoPC being associated with future risk of cardiovascular disease and visceral obesity[19, 26, 27]. In those cases, and for future studies it needs to be understood that these lipids can alter during storage even at low temperatures, and further work is required to understand the mechanistic basis of this and properly interpret cohort findings in light of this information.

We compared our data with the Bruneck cohort[19]. Findings there were similar to us (reduced lysoPC associated with CVD risk), but that study looked at prospective risk of an event, not genetic risk. In that case, most lysoPCs (unlike pretty much all other lipid classes they looked at) were inversely associated with CVD risk but only 3 achieved nominal statistical significance of $p<0.05$. None maintained significance after adjustment for multiple testing by means of the Benjamini-Hochberg procedure. However, using the three different selection procedures (Lasso, best subset, stepwise), LysoPCs were frequently selected and adding them to the lipid panel improved the C-index and Continuous net reclassification index (Manuel Mayr, personal communication). Thus, LPCs appeared to provide additional information compared to the other lipid classes that are mostly positively associated with CVD risk. This finding shows that a relatively small reduction in lysoPC is a consistent finding associated with risk. The observation that lysoPC measurement can add value to predictions of CVD risk strongly suggests a genetic component is involved and that they are pathophysiologically relevant even with a modest change. We propose that the genetic component may be accounted for by their association with a common CVD risk SNP present in up to 30% of the population.

The issue of sample processing, and storage reactions is a critical one that is often overlooked but a huge source of reproducibility problems when comparing lipidomics datasets across cohorts. Our study addresses this, and shows that it is essential to take this into account in order to properly interpret lipidomics datasets, and then to follow through to understand the underlying disease process. There are many examples of lipids that we know will be impacted by sample processing and storage but this is not often considered. We feel that this point is key and that our study highlights these issues to others who are working in this area. During preparation of this submission, we compared our processing pipeline with other studies, including Bruneck, where similar findings to ours were made relating to lysoPCs. In that case, samples were processed and frozen in the least possible time (Manuel Mayr, personal communication), and we believe their EDTA-anticoagulated plasma should be of a similar quality to that of the NPHSII cohort samples. Also, like ours, they were kept at -80, and had been in storage for around 10 yrs before analysis. We note that for UK Biobank, EDTA blood is first transferred overnight to a central processing centre, before plasma is then isolated and stored at low temperature. This suggests that significant care would be needed if measuring lysoPLs in this type of cohort, as there may be a large degree of variation related to time to freezing observed that could influence lysoPL and lysoPA levels.

**Supplementary Table 1:** MS parameters for analysing triacylglycerols by selected ion monitoring using the API 4000 (Sciex) platform

**Supplementary Table 2:** MS parameters for analysing free cholesterol and cholesterol esters by MS/MS using the API 4000 (Sciex) platform.

**Supplementary Table 3:** MS parameters for analysing lysoPLs by MS/MS using the API 4000 (Sciex) platform.

**Supplementary Figure 1. Comparison of fresh and cohort samples shows similar lipidomic results.** *Panel A. Total ion counts (TICs) of freshly-drawn plasma samples and controls from the Northwick Park Heart Study II (NPHSII) are similar.* Total ion current for each sample was integrated for the whole of the time of elution (n = 10 for old and new samples), and is shown as Tukey box plots. *Panel B. Integrated peak areas were compared for 19 lipids and are similar for fresh and cohort plasma.* Lipids were compared in fresh plasma or NPHSII samples (n = 10 for both) and are shown as Tukey box plots. * $p < 0.05$, ** $p < 0.01$, *** $p < 0.005$, 2-tailed, Mann Whitney U

**Supplementary Figure 2. CEs and TGs are similar between cohort and fresh plasma samples, but oxPL are significantly increased in storage.** *Panel A.* CEs were analysed in fresh human plasma and from NPHSII control samples by targeted lipidomics (n = 10 for both), shown as Tukey box plots. *Panel B*. TG molecular species were analysed in fresh human plasma and from NPHSII control samples by targeted lipidomics (n = 10 for both), shown as Tukey box plots. Panel C. oxPL were analysed in fresh human plasma and from NPHSII control samples by targeted lipidomics (n = 10 for both), shown as Tukey box plots. * $p < 0.05$, ** $p < 0.01$, *** $p < 0.005$, 2-tailed, Mann Whitney U

**Supplementary Figure 3. Integrated peak areas were compared for lysoPCs and were higher in cohort plasma.** Lipids were compared in fresh plasma or NPHSII samples (n = 10 for both) and are shown as Tukey box plots. * $p < 0.05$, ** $p < 0.01$, *** $p < 0.005$, 2-tailed, unpaired Student's T-test

**Supplementary Figure 4. Separate analysis of two sets of samples confirms reduced lysoPLs in GG samples versus AA**. *Panel A. LysoPLs were analysed using LC/MS/MS as in Methods in AA (39) and GG (33) samples.* The sample set used for global LipidFinder analysis was first compared using a targeted assay. *Panel B. Confirmation of decreased lysoPLs in a second sample set.* An additional set of samples of each genotype (AA: 47, GG: 49) were analysed using LC/MS/MS, as described in Methods. * $p < 0.05$, ** $p < 0.01$, *** $p < 0.005$, 2-tailed, unpaired Student's T-test (black) or Mann Whitney U (red)), shown as Tukey box plots.

**Supplementary Figure 5. Significant changes in lipid regulatory gene expression are observed with ANRIL knockdown in cell culture**. Volcano plots showing differential gene expression of all genes on the Affymetrix HuGene1.0 v1, chip. The horizontal dashed line shows where adj.pvalue < 0.05 (Benjamini-Hochberg correction) where points (genes) above this line are significantly differentially expressed. LysoPL regulating genes that alter in line with decreased levels of the lipids are labelled in black. Genes in red are annotated to the GO-term detailed in the plot title. Data are plotted in R using ggplot2. These volcano plots show additional GO terms that were significantly regulated in addition to those in Figure 5 of the main text.

**Supplementary Figure 6. Datasets for lysoPL metabolizing genes showing that removing the risk locus reverts gene expression back to levels in non-risk individuals**. * $p < 0.05$, ** $p < 0.01$, *** $p < 0.005$, Students t-test.

**Supplementary Figure 7. The profiles of TG or CE molecular species are not altered in the rs10757274 GG genotype**. TG or CE were measured using LC/MS/MS as outlined in Methods. Heatmaps of normalized mean values for AA and GG are shown.

**Supplementary Figure 8. Calculation of variance for assay performance, based on data obtained from Orbitrap data.** Boxplots of 15 abundant representative lipids detected in QC samples (n = 7). y-Linolenic acid, LPC(18:2), PI(36:1), PS(38:1), and CER(42:0) were identified in ESI negative mode and LPE(18:0), PC(36:2p)/(36:3e), PE(38:5), PA(38:2), PG(40:6), SMpe(43:0), SMpc(40:1), hexCER(36:2), DG(34:2) and TG(50:4), and %CV calculated, as shown on Tukey box plots, where box represents interquartile range and line represents median.

Supplementary Table 1

| Name | m/z Q1 | Name | m/z Q1 |
|---|---|---|---|
| TG(32:0) | 600,520 | TG(52:3) | 874,786 |
| TG(34:0) | 628,551 | TG(52:4) | 872,770 |
| TG(36:0) | 656,582 | TG(52:5) | 870,755 |
| TG(38:0) | 684,614 | TG(54:0) | 908,864 |
| TG(40:0) | 712,645 | TG(54:1) | 906,848 |
| TG(42:0) | 740,676 | TG(54:2) | 904,833 |
| TG(44:0) | 768,708 | TG(54:3) | 902,817 |
| TG(46:0) | 796,739 | TG(54:4) | 900,802 |
| TG(48:0) | 824,770 | TG(54:5) | 898,786 |
| TG(48:1) | 822,755 | TG(54:6) | 896,770 |
| TG(48:2) | 820,739 | TG(56:0) | 936,896 |
| TG(50:0) | 852,802 | TG(56:1) | 934,880 |
| TG(50:1) | 850,786 | TG(56:2) | 932,864 |
| TG(50:2) | 848,770 | TG(56:3) | 930,849 |
| TG(50:3) | 846,755 | TG(56:4) | 928,833 |
| TG(50:4) | 844,739 | TG(56:5) | 926,817 |
| TG(52:0) | 880,833 | TG(56:6) | 924,802 |
| TG(52:1) | 878,817 | TG (51:1)d5 | 869,837 |
| TG(52:2) | 876,802 | | |

Dwell time: 10 msec., declustering potential: 60 V, entrance potential: 10 V; all triacylglycerol species were analysed as $[M+NH_4]^+$ ions. Accordingly the Q1 m/z values refer to this ion species.

Supplementary Table 2

| Name | m/z Q1 | m/z Q3 |
|---|---|---|
| Cholesterol | 404,4 | 369,1 |
| CE(14:0) | 614,6 | 369,1 |
| CE(16:0) | 642,6 | 369,1 |
| CE(16:1) | 640,6 | 369,1 |
| CE(16:2) | 638, 6 | 369,1 |
| CE(18:0) | 670,6 | 369,1 |
| CE(18:1) | 668,6 | 369,1 |
| CE(18:2) | 666,6 | 369,1 |
| CE(18:3) | 664,6 | 369,1 |
| CE(20:3) | 692,6 | 369,1 |
| CE(20:4) | 690,6 | 369,1 |
| CE(20:5) | 688,6 | 369,1 |
| CE(22:6) | 714,6 | 369,1 |
| CE18:1 d7 | 675,6 | 376,3 |

Dwell time: 75 msec., declustering potential: 70 V, entrance potential: 10 V, collision energy: 20 V and collision cell exit potential: 25 V; all cholesterol species were analysed as $[M+NH_4]^+$ ions. Accordingly the Q1 m/z values refer to this ion species.

Supplementary Table 3

| Name | m/z Q1 | m/z Q3 |
|---|---|---|
| Lyso PC (16:0) | 480.3 | 255.2 |
| Lyso PC (16:1) | 478.3 | 253.2 |
| Lyso PC (18:0) | 508.4 | 283.3 |
| Lyso PC (18:1) | 506.4 | 281.2 |
| Lyso PC (18:2) | 504.3 | 279.2 |
| Lyso PC (20:1) | 534.4 | 309.3 |
| Lyso PC (20:3) | 530.3 | 305.3 |
| Lyso PC (20:4) | 528.3 | 303.2 |
| Lyso PE (16:0) | 452.3 | 255.2 |
| Lyso PE (16:1) | 450.3 | 253.2 |
| Lyso PE (18:0) | 480.3 | 283.3 |
| Lyso PE (18:1) | 478.3 | 281.2 |
| Lyso PE (20:4) | 500.3 | 303.2 |
| Lyso PE (22:6) | 524.3 | 327.2 |
| Lyso PE (18:2) | 476.3 | 279.2 |
| Lyso PC 18: d7 | 513.8 | 288.3 |
| Lyso PE 18:1 d7 | 485.6 | 288.3 |

Dwell time: 100 msec., entrance potential: 10 V, collision energy: 36 V, collision cell exit potential: 6 V. Declustering potential was 165 V and 112 V for lyso PC and lyso PE species, respectively. Lyso PC and Lyso PE species were detected as $[M-15]^-$ and $[M-H]^-$ ions, respectively.

Supplementary Figure 1

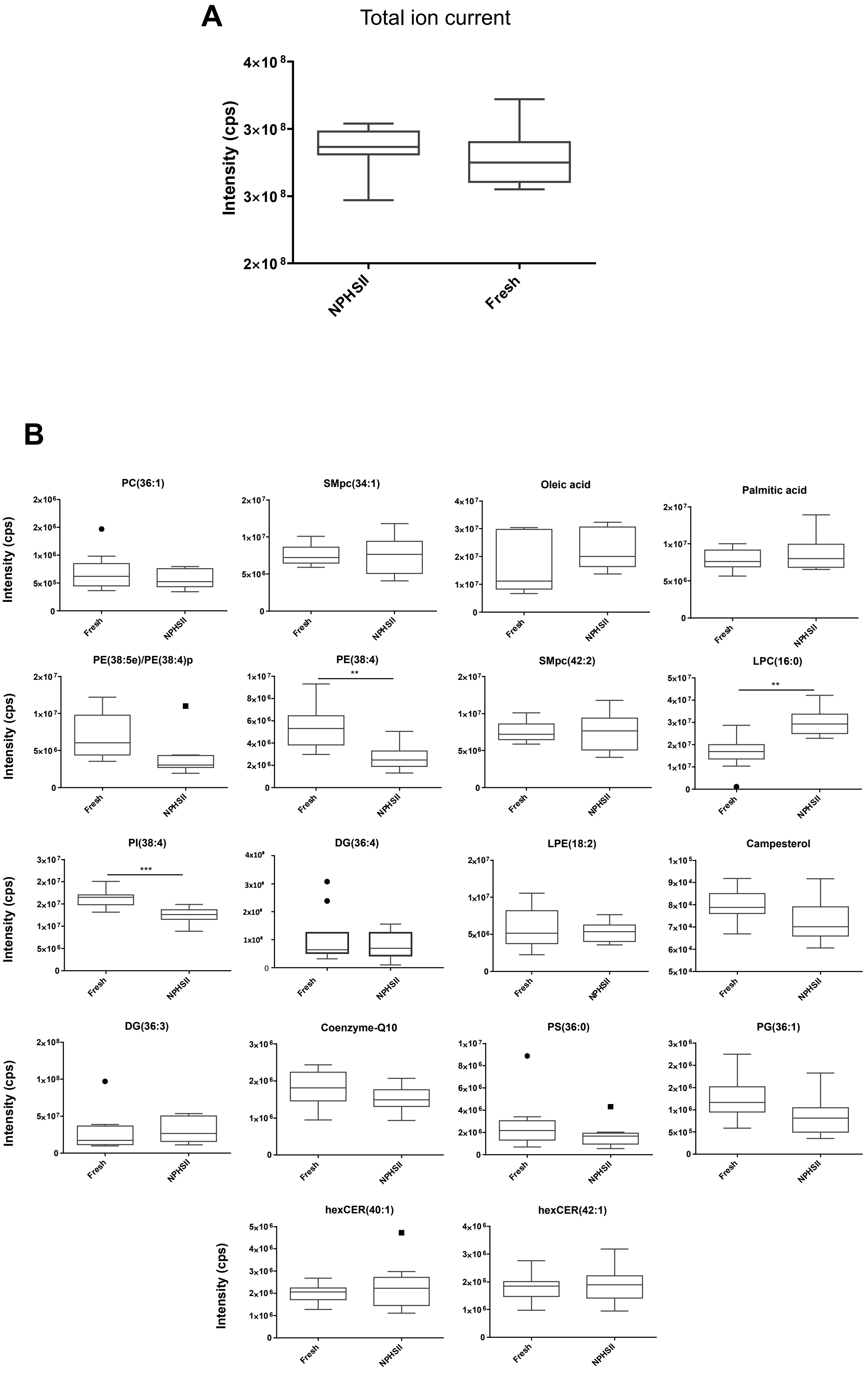

Supplementary Figure 2

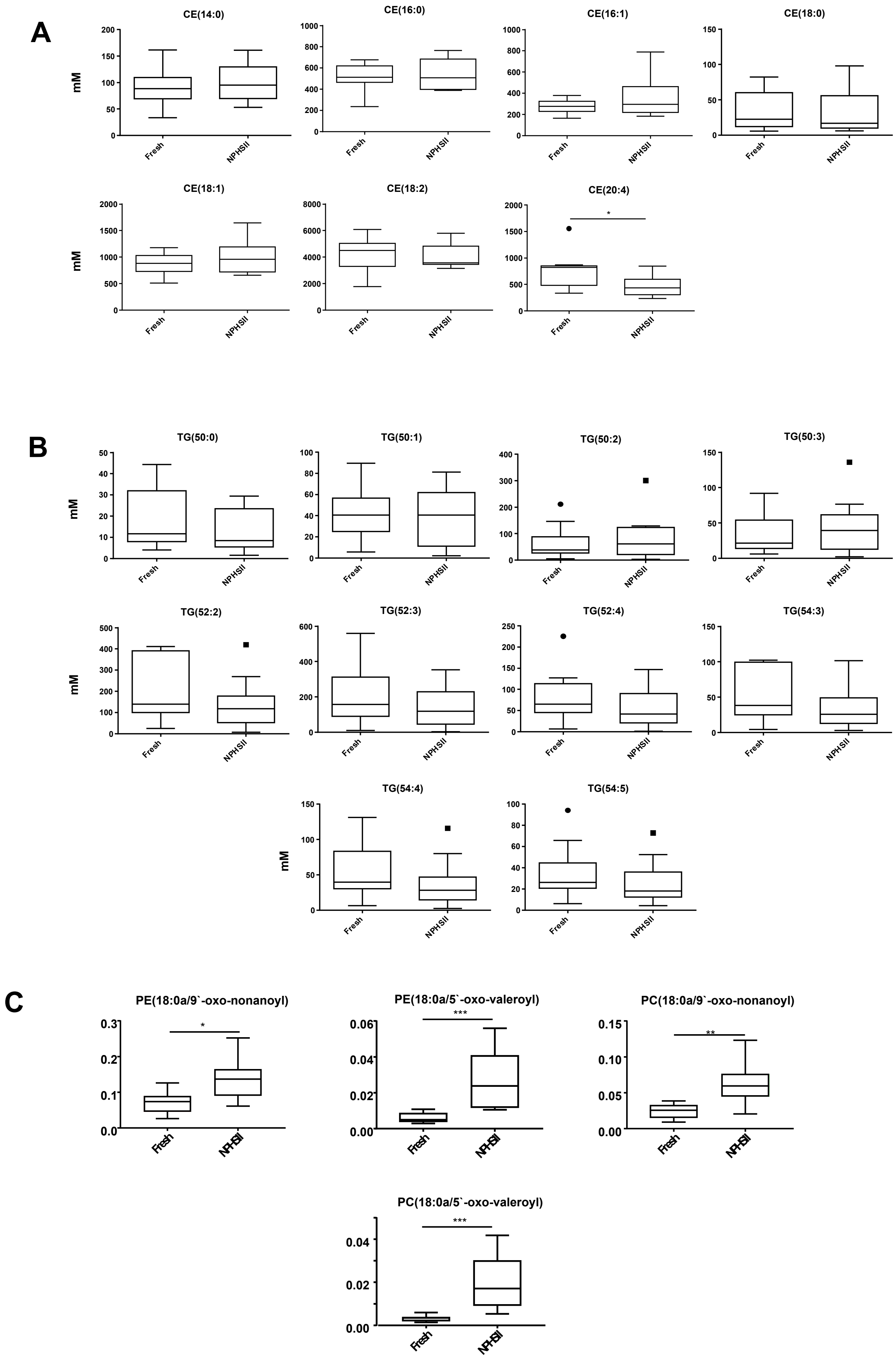

Supplementary Figure 3

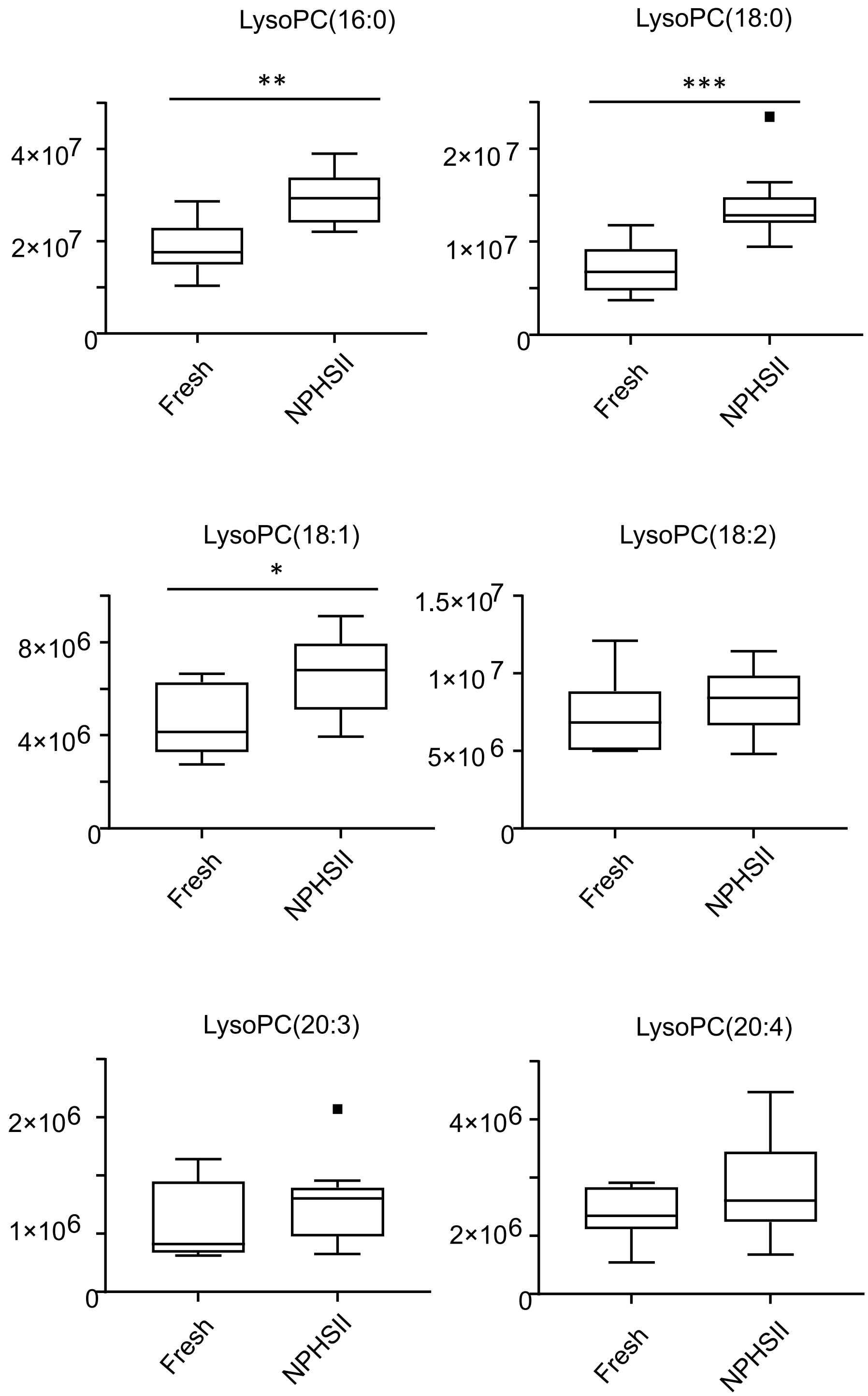

Supplementary Figure 4

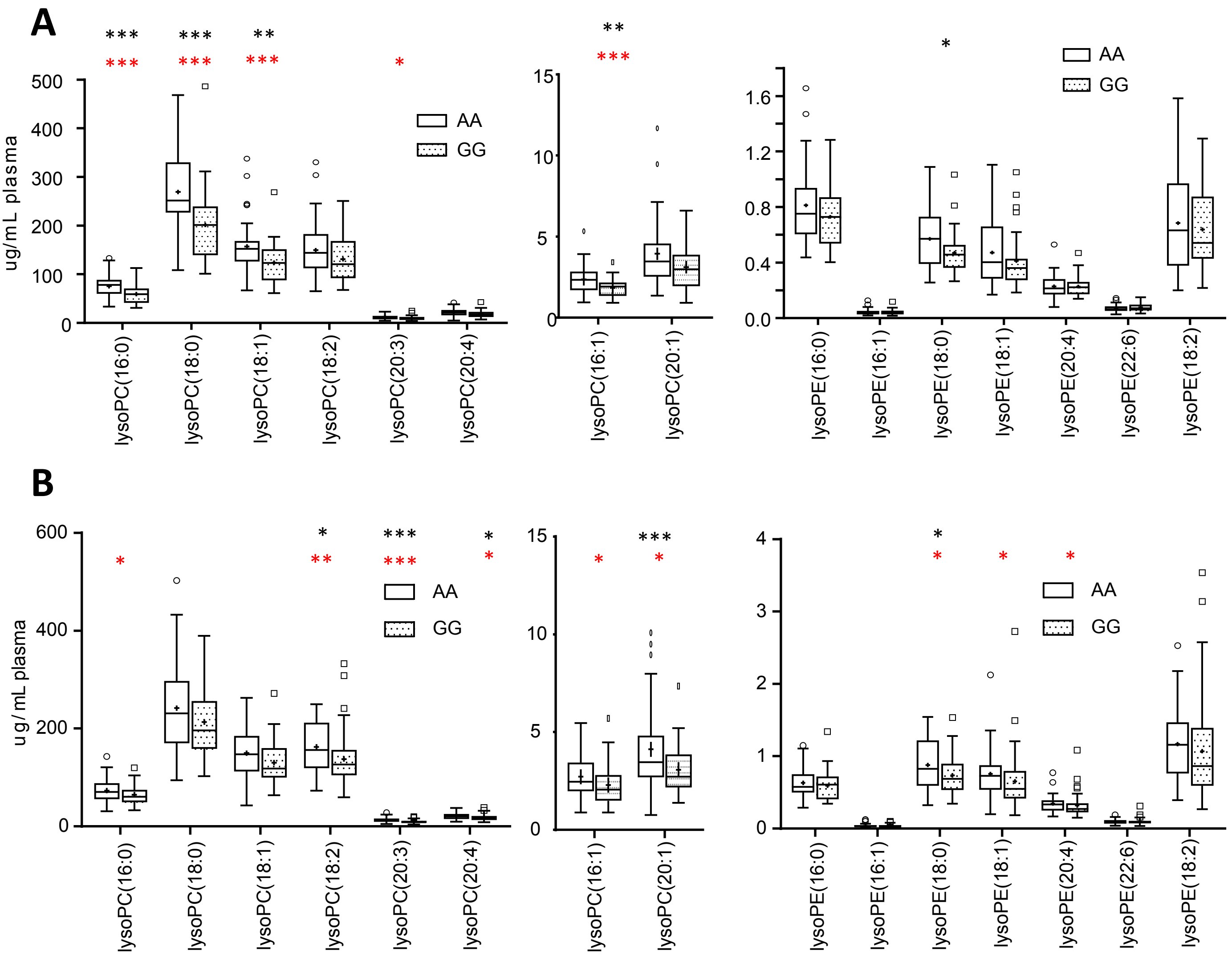

Supplementary Figure 5

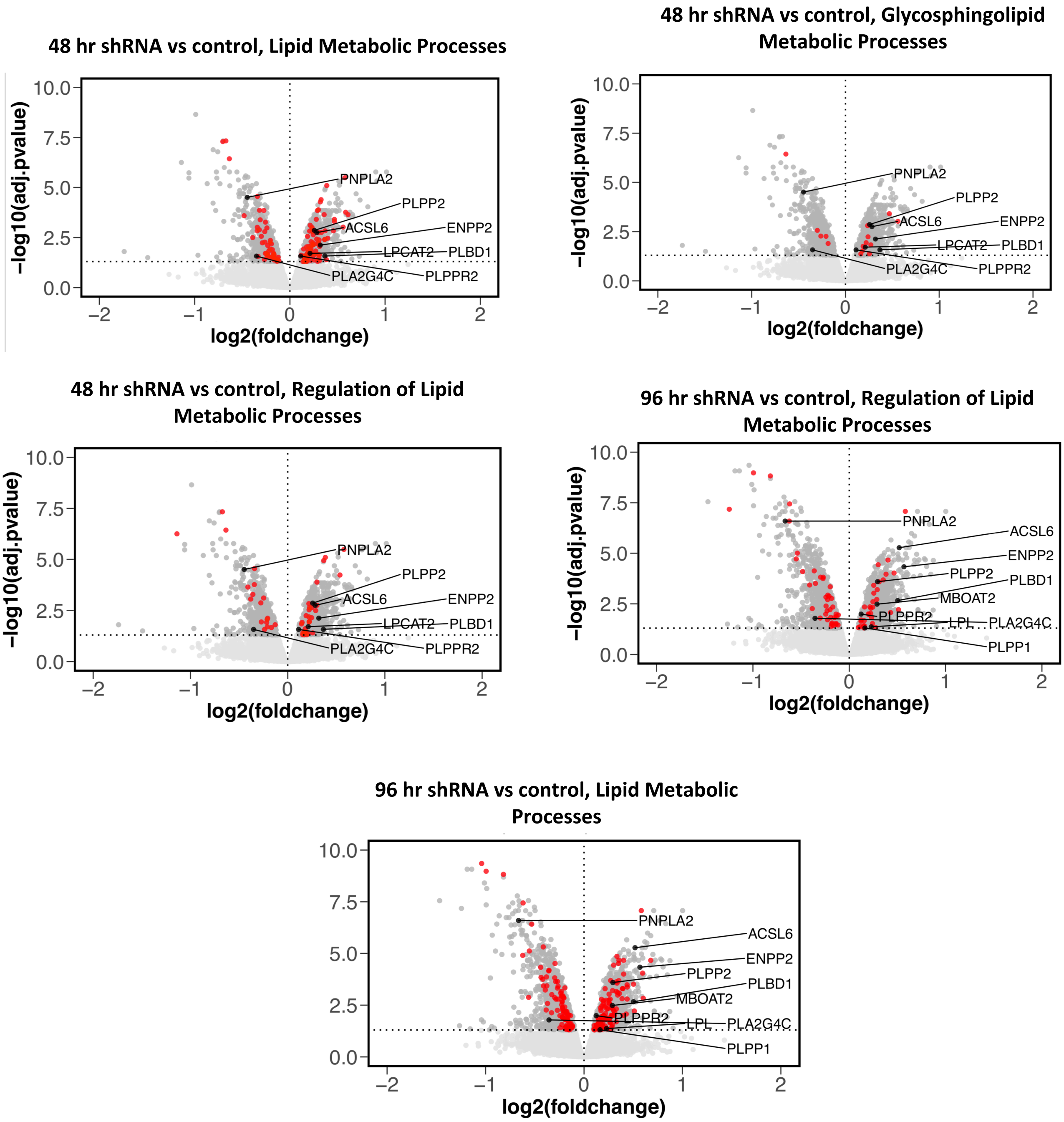

Supplementary Figure 6

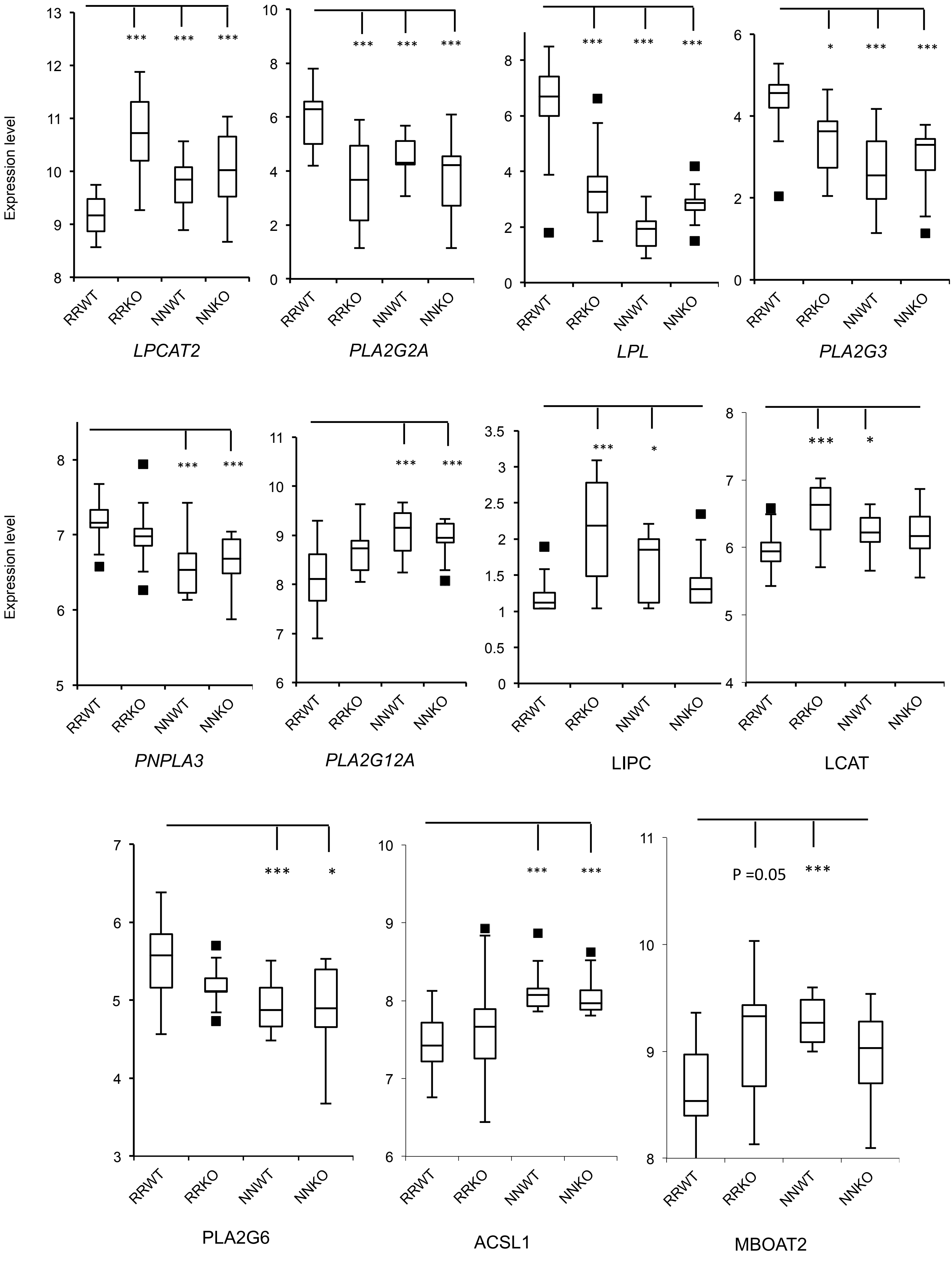

Supplementary Figure 7

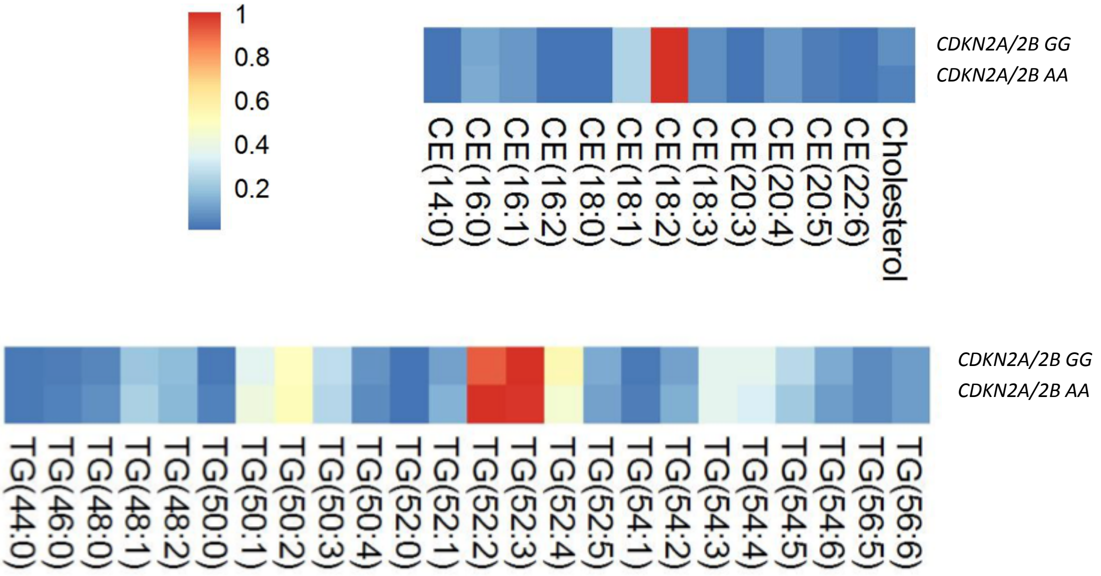

Supplementary Figure 8

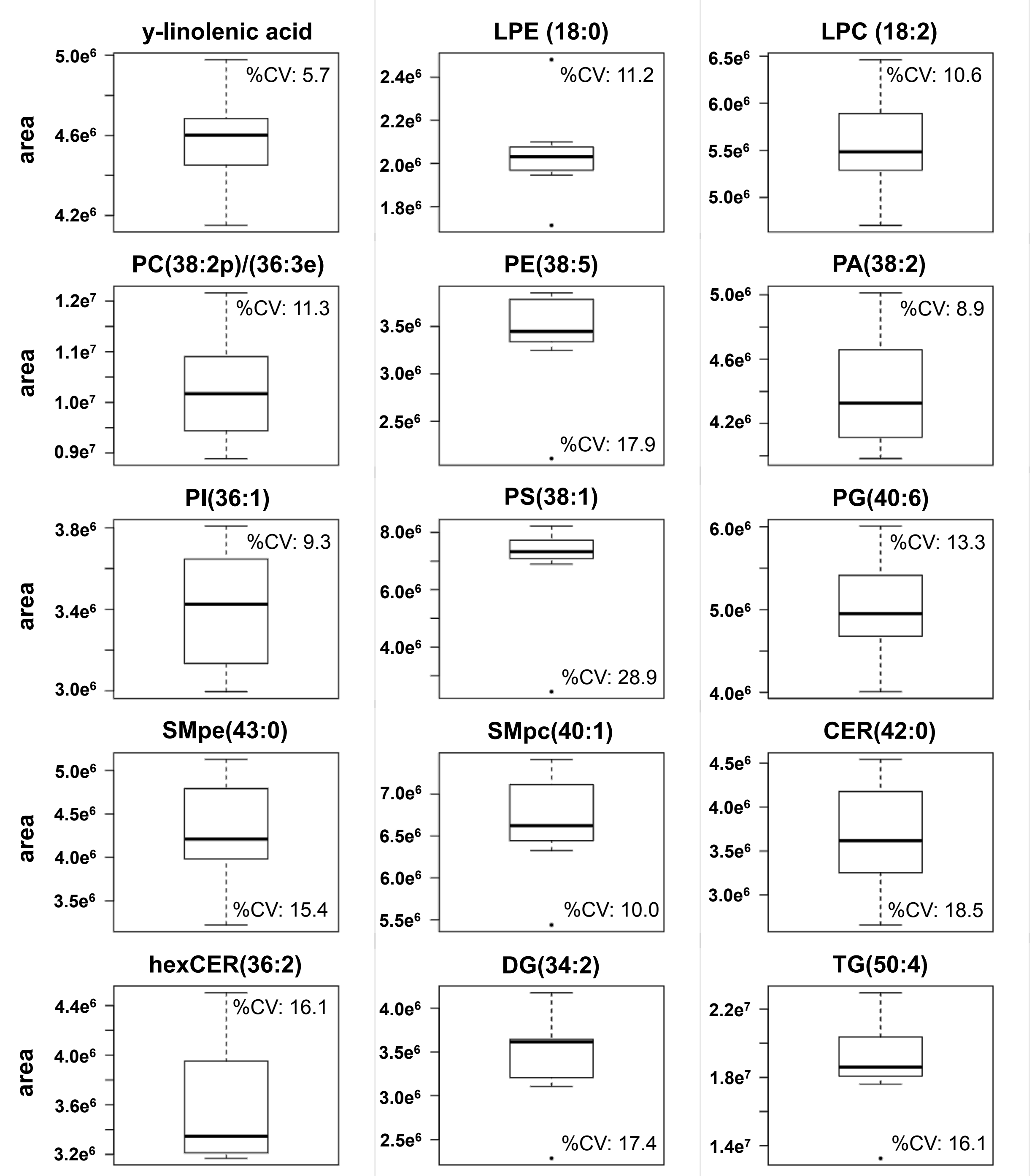
